# Epithelial uptake of *Aspergillus fumigatus* drives efficient fungal clearance *in vivo* and is aberrant in Chronic Obstructive Pulmonary Disease (COPD)

**DOI:** 10.1101/2022.02.01.478664

**Authors:** M. Bertuzzi, G.J. Howell, D.D. Thomson, R. Fortune-Grant, A. Möslinger, P. Dancer, N. Van Rhijn, N. Motsi, X. Du, A. Codling, R. Sash, M. Demirbag, E.M. Bignell

**Author notes:** **CORRESPONDING AUTHOR** Dr Margherita Bertuzzi, Manchester Fungal Infection Group, Faculty of Biology, Medicine and Health, The University of Manchester, Manchester Academic Health Science Centre, Core Technology Facility, Grafton Street, Manchester M13 9NT, UK. MRC Centre for Medical Mycology, University of Exeter, Geoffrey Pope Building, Stocker Road, Exeter EX4 4QD, UK. East Lancashire Hospitals NHS Trust, Lancashire, BB2 3HH, UK.

## Abstract

Hundreds of spores of the common mould *<u>A</u>spergillus <u>f</u>umigatus (Af)* are inhaled daily by human beings, representing a constant, often fatal, threat to our respiratory health. The small size of *Af* spores suggest that interactions with <u>A</u>irway <u>E</u>pithelial <u>C</u>ells (AECs) are frequent and we and others have previously demonstrated that AECs are able to internalise *Af* spores. We thus hypothesised that *Af* spore uptake and killing by AECs is important for driving efficient fungal clearance *in vivo* and that defective spore uptake and killing would represent major risk factors for *Aspergillus*-related diseases. In order to test this, we utilised single-cell approaches based on <u>I</u>maging <u>F</u>low <u>C</u>ytometry (IFC) and live-cell microfluidic imaging to measure spore uptake and outcomes *in vitro*, *in vivo* and using primary human AECs. *In vitro*, viability of immortalised AECs was largely unaffected by *Af* uptake and AECs were able to significantly curtail the growth of internalised spores. Applying our approach directly to infected mouse lungs we demonstrated, for the first time, that *Af* spores are internalised and killed by AECs during whole animal infection, whereby only ~3% of internalised spores remained viable after 8 hours of co-incubation with murine AECs. Finally, *in vitro* analysis of primary human AECs from healthy and at-risk donors revealed significant alterations in the uptake and consequent outcomes in Chronic Obstructive Pulmonary Disease (COPD), whereby gorging COPD-derived AECs were unable to quell intracellular *Af* as efficiently as healthy primary AECs. We have thus demonstrated that AECs efficiently kill *Af* spores upon uptake *in vivo* and that this process is altered in COPD, a well-known risk factor for debilitating fungal lung disease, thereby suggesting that AECs critically contribute to the efficient clearance of inhaled *Af* spores and that dysregulation of curative AEC responses represents a potent driver of *Aspergillus*-related diseases.

## Introduction

The mould pathogen *<u>A</u>spergillus <u>f</u>umigatus* (*Af*) causes a broad range of human diseases, whose individual manifestations and mortalities are entirely depending on the host immune status. <u>I</u>nvasive <u>A</u>spergillosis (IA) is associated with the highest mortality rates and causes an estimated 300,000 death per annum [1, 2], among which 10% represents patients with acute leukaemia, and recipients of allogenic hematopoietic stem cells and solid organ [3–5] and 3.9% patients with <u>C</u>hronic <u>O</u>bstructive <u>P</u>ulmonary <u>D</u>isease (COPD) [6–8]. <u>A</u>llergic <u>B</u>roncho<u>P</u>ulmonary <u>A</u>spergillosis (ABPA) is affecting more than 4 million asthmatic and cystic fibrosis sufferers [9, 10]. <u>C</u>hronic <u>P</u>ulmonary <u>A</u>spergillosis (CPA) exacerbates pre-existing structural and immunological lung defects and it is estimated to affect more than 3 million of people worldwide, with 50% mortality in the first 5 years after diagnosis [11, 12].

From the allergic hyper acute response observed in ABPA sufferers to the most rapidly progressing IA in severely immunocompromised patients, all disease manifestations are unified by a common pattern of infection, which initiates with the inhalation of fungal conidia and eventually leads to the destruction of the pulmonary parenchyma. *Af* conidia are abundant in the airborne microflora [13] and have the ideal diameter (2-3 μm) for deep deposition within the bronchial tree and alveoli [14]. In healthy individuals, inhaled conidia are efficiently cleared by the concerted action of the innate immune system, mostly macrophages and neutrophils [15–17]. However, <u>A</u>lveolar <u>M</u>acrophages (AM) only constitute ~ 3.25% of the total cell number in the alveoli [18] and therefore are unlikely to be the first cell type encountered by the inhaled spores. Furthermore, macrophage ablation in mice using clodronate [17] and *in silico* modelling of infection [19, 20], indicate that AM migrate too slowly and randomly in the absence of a chemoattractant signal and are unable to protect against *Af* infection alone. Also, *in silico* [19] and murine studies [17, 21] indicate late recruitment of neutrophils to the site of infection. On the contrary, the contact of <u>A</u>irway <u>E</u>pithelial <u>C</u>ells (AECs) with *Af* conidia is instant, extensive, and likely prolonged and recent *in silico* modelling [20, 22, 23] and *in vitro* studies [24, 25] suggest that AECs play a key role in initiating fungal clearance and host responses, by releasing cytokines for immune cell recruitment during infection.

We previously exploited an *in vitro* alveolar infection assays to address the mechanistic basis of pathogen-mediated tissue damage, finding that *Af* causes epithelial decay in a multiphasic fashion involving an initial contact-dependent mechanism followed by damage caused by soluble effectors [25, 26]. Non-invasive mutants lacking the transcription factor PacC are deficient in both of these processes and consequently attenuated for virulence in a leukopenic model of infection [26]. Furthermore, nystatin protection assays indicated that Δ*pacC* mutants are less present inside AECs compared to the respective parental isolates [26] and that, together with the attenuation of the mutants during murine infection, lead us to hypothesise that contact-dependent perturbations of epithelial integrity might in part result from uptake of fungal spores. We and other reported that *Af* spores are internalized by AECs; for example, *in vitro,* A549 alveolar type-II like cells [26–30] and 16HBE14o-transformed human bronchial epithelial cells [31, 32] internalize 30-50% of the spores they come into contact with. Pre-treatment of A549 monolayers with monoclonal anti-Dectin-1 antibody and the endocytosis inhibitor Cytochalasin D in the nystatin protection assay demostrates that uptake of *Af* contacting alveolar epithelia occurs in a cell wall-, actin- and Dectin-1 dependent manner [26–28]. Popolation-scale analyses of *Af*-AECs interactions *in vitro* indicate that the majority of the internalized spores are killed, but a small proportion (3%) survives and germinates inside acidic organelles [29], thereby suggesting that AECs might serve as a fungal reservoir for latent occupation and immune evasion (as reviewed in [33–35]). Indeed, the presence of Cytochalasin D in *in vitro* infection assay conveys, to some extent, protection of monolayer integrity during co-incubation with wild-type *Af* isolates [26] supporting an important role for *Af* spore uptake during invasion of the pulmonary epithelium and consequent damage. However, Chaudhary *et al.*, 2012 reported that bronchial epithelial cells carrying a non-functional mutated version of the Cystic Fibrosis Transmembrane conductance Regulator (CFTR, DF508) are impaired in the uptake and killing of conidia, thereby strongly supporting a curative role for epithelial activities [36].

The relevance of AEC-mediated activities, such as spore uptake, to disease outcome remains unclear; nevertheless, recent research spanning bacterial and fungal respiratory pathogens [37] has demonstrated that the respiratory epithelium orchestrates a multifaceted and active response to the presence of inhaled pathogens. AECs ability to act as non-professional phagocytes, therefore contributing to clearance of respiratory pathogen load, has only been recently recognised (as reviewed in[37]); as yet, it is poorly understood mechanistically and in the context of human disease, especially of fungal aetiology. On the other hand, many respiratory pathogens invade frequently non-phagocytic host cells (epithelial, but also endothelial cells) to cause disease. Invasion of epithelial cells might benefit the pathogen by providing not only a way to traverse a natural cellular barrier but also a shield from professional phagocytes as macrophages and neutrophils, and a nutrient enriched intracellular niche.

The focus of this study was therefore to understand the role of spore uptake in disease outcome by utilising direct single-cell approaches based on Imaging Flow Cytometry (IFC) [38] and live-cell microfluidic imaging to identify, quantify and isolate individual host pathogen complexes from AECs infected with *Af*. Using these approaches, we were able to establish that *Af* spores are internalized by immortalized and primary human AECs, rapidly and in a time-dependent manner leading to significant fungal killing with minimal impact on epithelial viability. Furthermore, applying the same IFC single-cell approach to infected and dissociated mouse lungs we demonstrated, for the first time in whole animals, that *Af* spores are internalized and killed by type-II alveolar epithelial cells. Importantly, using primary human AECs from healthy and COPD donors, we also demonstrated that lung disease significantly impact *Af* uptake. Both commercially available and locally sourced COPD-derived AECs were found to internalize significantly more *Af* spores but to lack the ability to quell the intracellular fungus as efficiently as their healthy counterparts, thereby indicating that aberrancies in the uptake process might lead to heightened susceptibility to fungal infection. The integration of the data obtained demonstrates that uptake and killing of fungal spores by immortalized and primary murine and human AECs can provide a potent means of control of inhaled *Af* spores, hence substantiating the importance of understanding the mechanistic basis of these processes for the identification of immunomodulators to facilitate treatment and limit respiratory damage.

## Results

### Spore uptake by the airway epithelium delays intracellular *Af* germination

To assess the timing of *Af*-AECs interactions and the rate of spore internalisation by cultured airway epithelia, the *Af* isolate A1160^+^ [39] was genetically engineered to express the highly photo-stable and bright red fluorescent protein tdTomato [40] **(Fig. S1)**. A549 monolayers were incubated for 2, 4 and 6 hours with the tdTomato-fluorescent *Af* isolate in the A1160^+^ genetic background, named A1160^+/tdT^. Differential fluorescence staining of extracellular *Af* was performed using the cell impermeant fluorescent stain Calcofluor White, which binds to the fungal cell-wall component chitin **(Fig. 1A)**. Epithelial and fungal cell surfaces were stained using ConA-FITC, a lectin binding protein conjugated with the green fluorescent dye fluorescein isothiocyanate (FITC). Fluorescence microscopy analysis of *Af* uptake by A549 cells demonstrated that this process was time-dependent, occurring from very early time-points (2 hours) of co-incubation **(Figs. 1B and C)**. Enumeration of the intra- and extracellular A1160^+/tdT^ was used to calculate the uptake index, expressing the percentage of internalized A1160^+/tdT^ on the total number of observed fungal elements, including intracellular and extracellular ones. Comparison of uptake indexes at different time points of the *Af*-AECs interaction across biological triplicates revealed a time-dependent increase in the number of internalized spores (relative to total pathogen inoculum) from 17% at 2 hours to 64% at 6 hours post-infection **(Fig. 1B)**. Fluorescence microscopy showed that single AECs were able to internalize multiple *Af* spores and remarkably, internalized *Af* was delayed for germinative growth relative to the extracellular counterparts **(Fig. 1C)**. These findings unmasked the complex and multifaceted nature of *Af*-AECs interactions and highlighted the necessity to develop state-of-the-art high-throughput means of investigating these interactions at the single-cell level.

**Fig. 1:**
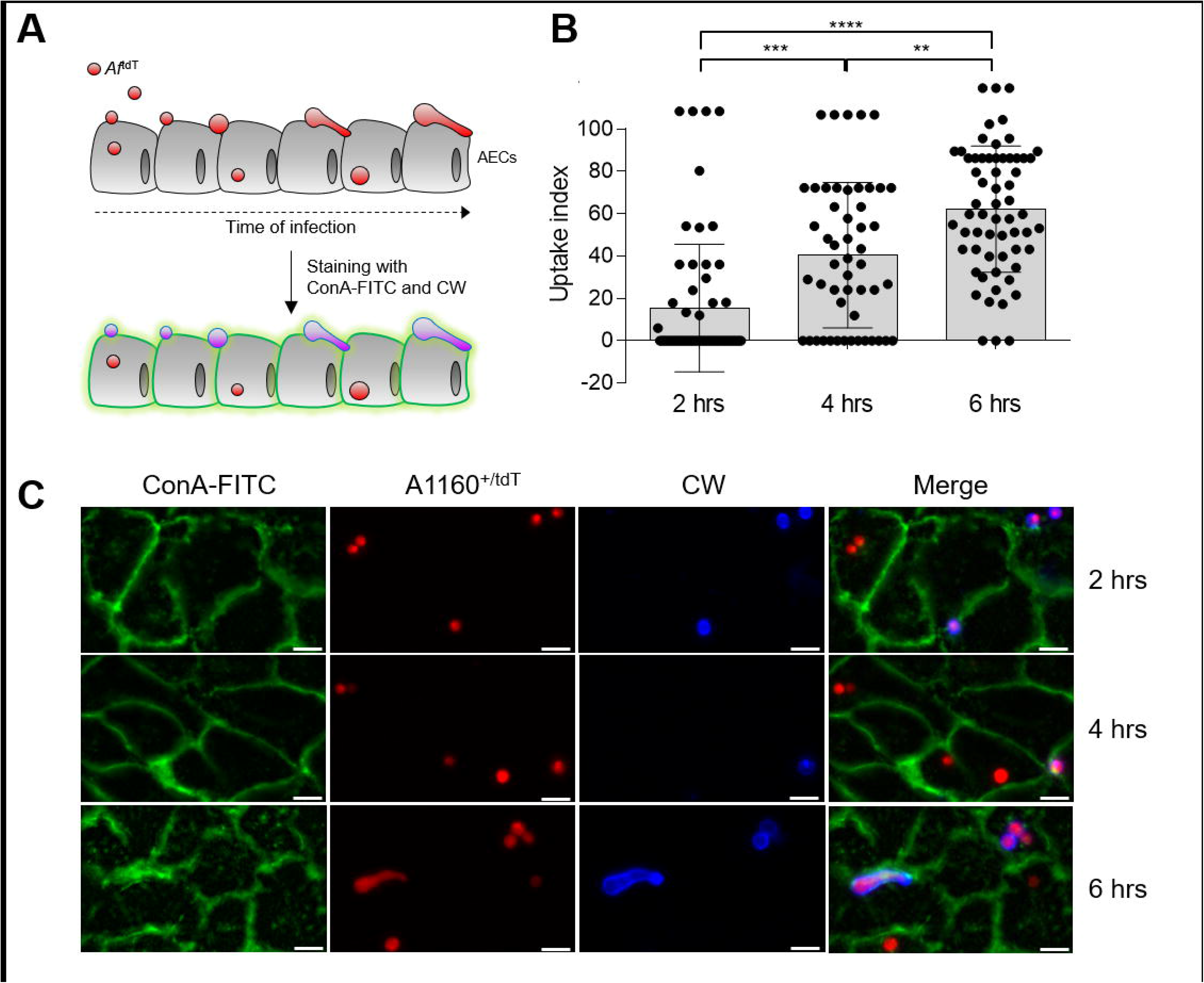
Differential fluorescence microscopy of A1160^+/tdT^ interacting with A549 cells shows delayed germination of intracellular *A. fumigatus (Af).* **(A)** Principle of the assay. A549 monolayers are incubated with 10^5^ spores/ml of a tdTomato-fluorescent *Af* isolate (*Af*^tdT^) for 2-6 hrs. Labelling of extracellular spores with Calcofluor White (Blue) allows to distinguish intracellular (Red) and extracellular (Red + Blue = Purple) populations. Fungal and mammalian cell surface is labeled using ConA-FITC (Green). **(B)** Quantification of spore internalisation following infection of A549 monolayers with 10^5^ spores/mL of the strain A1160^+/tdT^ for 2, 4 and 6 hours (n ≥ 31 fields of view among biological duplicates, non-parametric Kruskal-Wallis test with Dunn’s multiple comparisons test). (C) Representative images of A549 monolayers infected with 10^5^ spores/mL of A1160^+/tdT^ for 2, 4 and 6 hours. ****p ≤ 0.0001, ***p ≤ 0.001, **p ≤ 0.01, and *p ≤ 0.05

### A novel single-cell method to survey *Af*-AECs interactions allows the high-throughput parameterization of *Af* uptake *in vitro*

In order to perform high-throughput parameterization of rate, stoichiometry and outcome of *Af* uptake by A549 alveolar epithelia, we combined the differential fluorescence staining of tdTomato-expressing *Af* stains, such as A1160^+/tdT^, via Calcofluor White with state-of-art Imaging Flow Cytometry (IFC) [38]. In *in vitro* infection assays, the multiplex assay developed allows to visualize, quantify and analyse at the single-cell level i) AECs not in contact with *Af* (AEC_n_), ii) AECs which have internalized *Af* spores (AEC_i_) and iii) AECs which are attached to fungal elements (AEC_a_) **(Fig. 2A)**. In agreement with the data obtained using fluorescence miscroscopy **(Fig. 1)**, IFC enumeration of AECs internalizing the A1160^+/tdT^ isolate from 2 to 6 hours post-infection showed a rapid, significant and sustained spore uptake during *in vitro* infection, whereby the number of AEC_i_ at 6 hours of infection (n = 34.9 ± 6.9 cells on 2000 cells surveyed) was 4 times higher than at 2 hours (n = 9.4 ± 4.5 cells on 2000 cells surveyed) **(Fig. 2B)**. Visual examination of the IFC-derived fluorescence microscopy panels confirmed that single AECs are able to internalize multiple *Af* spores **(Fig. 2A)**. While most of the *Af* elements attached onto AEC_a_ had germinated within 8 hours of infection, most of the *Af* spores internalized by AEC_i_ were ungerminated or delayed for germinative growth in comparison to their extracellular counterparts **(Fig. 2C)**. We therefore measured the percentage of *Af* germination in AEC_a_ (n = 171) and AEC_i_ (n = 696) after infection of A549 cells for 8 hours and indeed determined that, while 68.4% (± 15.59%) of the AEC_a_ were attached to germinated *Af,* only 14.5% (± 4.2%) of the AEC_i_ were containing germinated spores **(Fig. 2D)**. Collectively, the results obtained using both fluorescence microscopy and IFC suggested that *Af* might undergo killing upon uptake by AECs and that differential uptake and/or killing of *A. fumigatus* mutants, such as the Δ*pacC* ones, by AECs might contribute to the attenuation of pathogenicity observed *in vivo* during infection in the leukopenic murine model of infection [26].

**Fig. 2:**
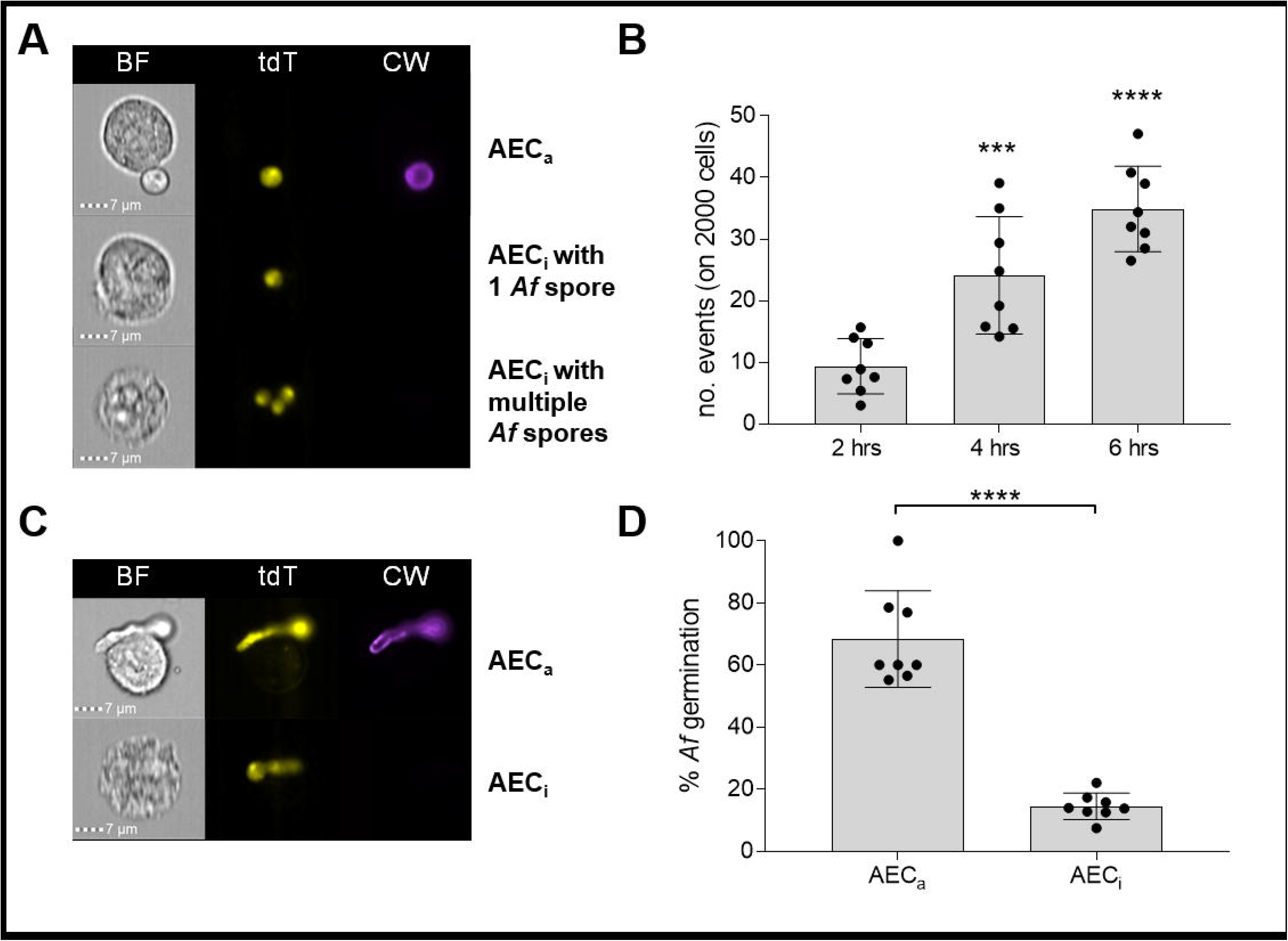
Single-cell analysis of *Af-*AECs interactions via IFC allows quantitation of spore uptake and germination during *in vitro* infections. Infection of A549 monolayers with 10^5^ spores/mL of the strain A1160^+/tdT^. **(A)** IFC exemplary panels after infection for 4 hr showing AECs which have *Af* on their surface (AEC_a_) or inside (AEC_i_). A single AEC_i_ can contain multiple spores. **(B)** Quantification of AEC_i_ after infection for the indicated time points. Number of AEC_i_ is derived on 2000 total cell number analysed (biological triplicates with technical duplicates/triplicates, non-parametric Kruskal-Wallis test with Dunn’s multiple comparisons test relative to uninfected control). **(C)** IFC exemplary panels after infection for 8 hr showing differential germination for *Af* on or inside AEC_i_. **(D)** Percentage of *Af* germination after infection for 8 hr (biological triplicates with technical duplicates/triplicates, unpaired t-test). ****p ≤ 0.0001, ***p ≤ 0.001, **p ≤ 0.01, and *p ≤ 0.05

### The attenuated Δ*pacC* mutant is more efficiently killed upon internalization by AECs

Indirect quantification of spore uptake using a nystatin protection assay previously indicated that, upon infection of A549 monolayers, viable counts of intracellular *Af* Δ*pacC* mutants were reduced by half relative to the respective parental isolates [26]. However, from the output of the nystatin assay is it not possible to unequivocally establish if, in comparison with the respective parental isolates, Δ*pacC* mutants are less efficienty internalized by AECs, more efficiently killed once inside AECs or a combination of both. To analyse the rate, stoichiometry and outcome of *Af* Δ*pacC* mutant uptake by alveolar epithelia, the *Af* Δ*pacC* mutant in the A1160^+^ genetic background, namely Δ*pacC*^A1160+^ [41], was genetically engineered to express tdTomato [40] **(Fig. S1)**. Mutants lacking conidial dihydroxynaphthalene (DHN) melanin were also previously found to be less efficiently internalized by, and more efficiently killed within AECs [42] and to display attenuated virulence in murine infection models[43, 44]. Therefore, as control, an *Af* Δ*pksP* mutant [41] in the same genetic background was also genetically engineered to express the same fluorescent protein **(Fig. S1)**. Infections of the A549 monolayers with Δ*pacC*^A1160+/tdT^, Δ*pskP*^A1160+/tdT^ and A1160^+/tdT^ were carried out for 6 hours. IFC-comparison of uptake rates showed only a slight decrease in AEC_i_ number for Δ*pacC*^A1160+/tdT^ (865.9 ± 96.1) and Δ*pskP*^A1160+/tdT^ (857.9 ± 132.5) infections relative to A1160^+/tdT^ ones (998.0 ± 84.1) **(Fig. 3A)**. This slight decrease was only marginally recapitulating our previous results obtained with the nystatin protection assay and demonstrating a drastic reduction in the recovery of intracellular Δ*pacC* spores from AECs compared to their respective parental counterpart [26]. Therefore, the number of A1160^+/tdT^, Δ*pacC*^A1160+/tdT^ and Δ*pskP*^A1160+/tdT^ fungal elements inside each AEC_i_ was enumerated to determine the stoichiometry of uptake of the strains. The number of AEC_i_ containing more than one spore of A1160^+/tdT^ (27.5 ± 3.4%) was significantly higher than for Δ*pacC*^A1160+/tdT^ (19.7 ± 3.5%) **(Fig. 3B)**. Consequently, the fungal burden within all AEC_i_ surverved was significantly higher in the A1160^+/tdT^ infections than in the Δ*pacC*^A1160+/tdT^ counterparts, with 1473 (± 174.0) A1160^+/tdT^ spores, but only 1112 (± 121.8) Δ*pacC*^A1160+/tdT^ and 1255 (± 192.6) Δ*pskP*^A1160+/tdT^ spores internalized among 8000 AECs analysed **(Fig. 3C)**. This findings indicated that, relative to the parental isolate and in agreement with the nystatin protection assay, the non-invasive and attenuated Δ*pacC* mutant was less efficiently internalized by AECs.

**Fig. 3.**
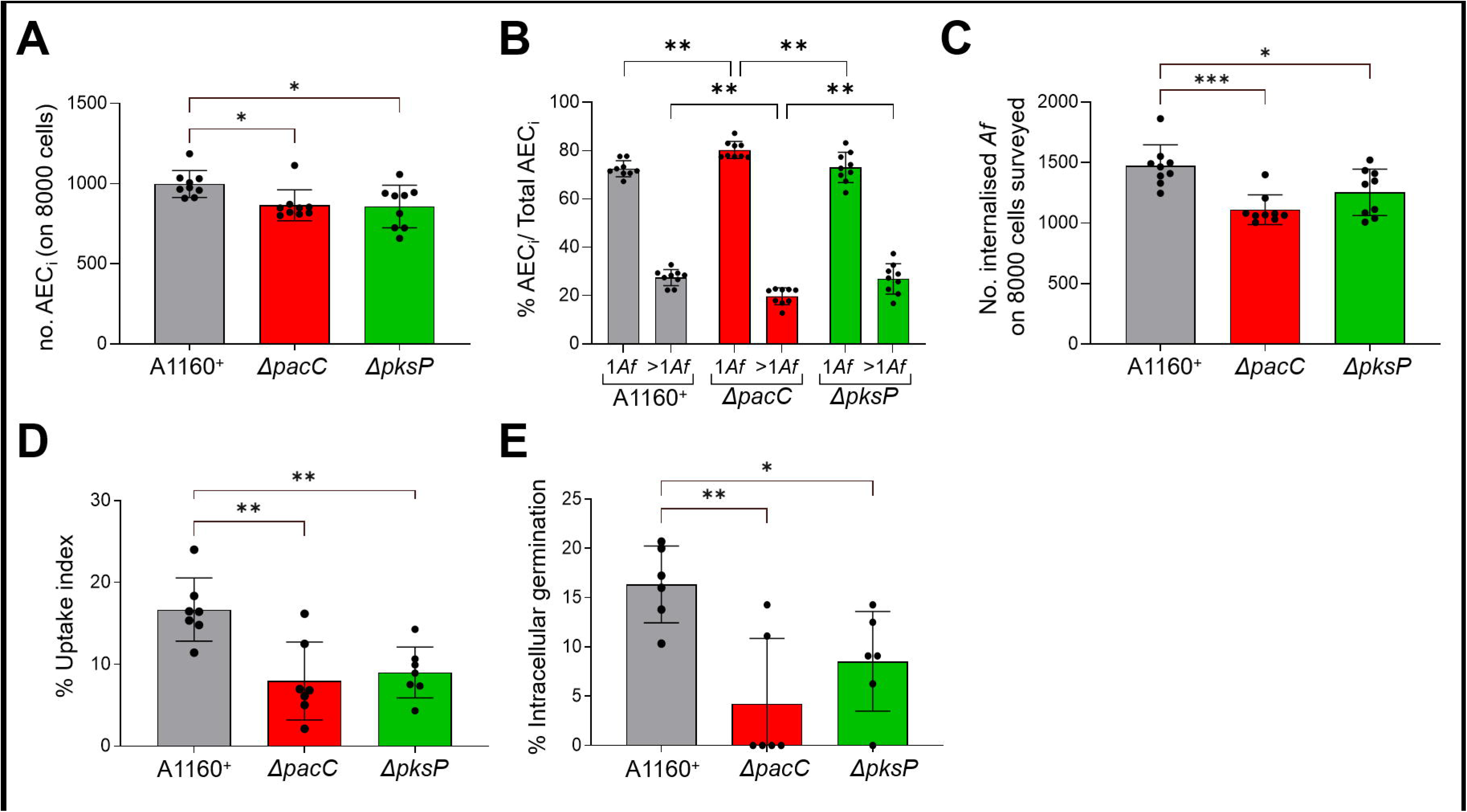
Single-cell comparison of uptake rate and stoichiometry shows that both Δ*pacC*^A1160+/tdT^ and Δ*pksP*^A1160+/tdT^ are less avidly internalised by AECs than the parental isolate A1160^+/tdT^. **(A)** Quantification of AEC_i_ on 8000 cells surveyed by IFC. **(B)** Percentage of AEC_i_ containing one or more *Af* spore relative to the total number of AEC_i_ analysed by IFC. **(C)** Number of *Af* spores internalised on 8000 AEC surveyed in A by IFC. **(D)** Uptake index and **(E)** percentage of intracellular germination as quantified by live-cell microfluidic imaging. For A-D, infections of A549 monolayers with *Af* strains were carried out for 6 hours, while in E infections were carried out for 20 hrs (in biological and technical triplicates, ordinary 1-way ANOVA with Holm-Sidak’s multiple comparisons). ****p ≤ 0.0001, ***p ≤ 0.001, **p ≤ 0.01, and *p ≤ 0.05

To determine whether differences in intracellular fate were also contributing to the drastic reduction in the recovery of intracellular Δ*pacC* spores (relative to the parental isolate) demonstrated in our nystatin protection assay [26], a microfluidic infection system was developed to perform live-cell semi-automated analyses of epithelial monolayers exposed to fungal challenge **(Figs. 3D-E)**. In support of our previous observations **(Fig. 3A)**, uptake index after 6 hours of infections with the Δ*pacC*^A1160+/tdT^ (8.0 ± 4.8) and Δ*pskP*^A1160+/tdT^ (9.0 ± 3.1) isolates was found to be significantly lower than that for A1160^+/tdT^ infections (16.7 ± 3.9) **(Fig. 3D)**, thereby confirming the suitability of this system to study *Af* uptake by AECs and more broadly *Af*-AECs interactions. The intracellular fate on internalized *Af* was then followed in real time for further 14 hrs and intracellular germination was quantified and compared between strains **(Fig. 3E)**. Intracellular germination of Δ*pacC*^A1160+/tdT^ (4.2 ± 6.6) and Δ*pskP*^A1160+/tdT^ (16.3 ± 3.9) isolates was significantly impaired compared to A1160^+/tdT^ isolates (16.3 ± 3.9) **(Fig. 3E)**. Taken together, these results indicate that attenuated *Af* mutant lacking *pacC* is less efficiently internalized by AECs than its parental counterpart **(Figs 3A-D)**; however, its germination is also prevented more efficiently than the parental one upon internalization by AECs **(Fig. 3E)**, suggesting the mutant might be more prone to intracellular killing. While the full extent of the attenuation of *Af* ∆*pacC* mutant likely extends beyond defective internalization and intracellular processing, this data suggest that internalization of *Af* spores, which is hampered in the absence of *pacC*, might contribute to the disaggregation of infected monolayers, as previously hyphotesised [26].

### *Af* spore uptake is not the major driver of AEC death upon *Af*-AEC interaction

*In vitro* infection assays have previously shown that *Af* causes the disaggregation of cultured A549 monolayers in a multiphasic fashion involving an initial contact-dependent mechanism and that Δ*pacC* mutants are defective with regards to this mode of pathogen-mediated tissue damage [26]. This lead us the the hyphothesis that contact-dependent perturbations of epithelial integrity might in part result from uptake of fungal spores, a process which is reduced for the Δ*pacC* mutants **(Figs. 3A-E)**, hence potentially explaining their attenuation of virulence *in vivo*. To survey the outcome of *Af*-AEC interactions, the optimized multiplex IFC assay was expanded to include Annexin-V-FITC (Anx-FITC) and TO-PRO™-3 (TO-PRO3) staining [38], which allow to quantify different modes of cell death, namely apoptosis and necrosis, respectively. Following infection of A549 monolayers with the A1160^+/tdT^ and Δ*pacC*^A1160+/tdT^ isolates for 16 hours, the number of live (Anx-FITC^−^/TO-PRO3^−^), necrotic (Anx-FITC^−^/TO-PRO3^+^), apoptotic (Anx-FITC^+^/TO-PRO3^−^) and late apoptotic (Anx-FITC^+^/TO-PRO3^+^) were enumerated and expressed as a percentage relative to the total number of AECs analysed **(Fig. 4A)**. When comparing with uninfected monolayers, a significant decrease (15.2% ± 11.2) of live AECs and congruently, a significant increase (13.8% ± 12.4) of necrotic AECs was measured upon A549 infection with the A1160^+/tdT^ isolate **(Fig. 4A)**. However, no such decrease in live AECs, neither an increase in necrotic AECs, was detected upon A549 infection with the Δ*pacC*^A1160+/tdT^ isolate compared to uninfected monolayers **(Fig. 4A)**. This is in agrement with our previously published results using *in vitro* infection assays to measure pathogen-mediated host-damage [26], whereby *Af* causes epithelial decay since early-stages of the *Af*-AEC interaction and the non-invasive Δ*pacC* mutants lack this ability [26]. To analyse the outcome of spore uptake with respect to host survival or cell death, the percentage of live AEC and AEC_i_ upon infection with the A1160^+/tdT^ and Δ*pacC*^A1160+/tdT^ isolates was compared to the uninfected counterpart. Surprisingly, AEC internalizing both the A1160^+/tdT^ and Δ*pacC*^A1160+/tdT^ isolates did not undergo more cell death than uninfected monolayers and at 16 hours post-infection, 94.0% of A1160^+/tdT^-containing AECs and 89.8% of the Δ*pacC*^A1160+/tdT^-containing AECs were still alive **(Fig. 4B)**. Collectively, our results indicate that, while AEC interaction with *Af* leads to a visible and measurable (by several means) disaggregation and damage of the infected monolayers [26, 38], *Af* spore uptake is not the main cause of the epithelial cell death detected upon *Af*-AEC interaction.

**Fig. 4:**
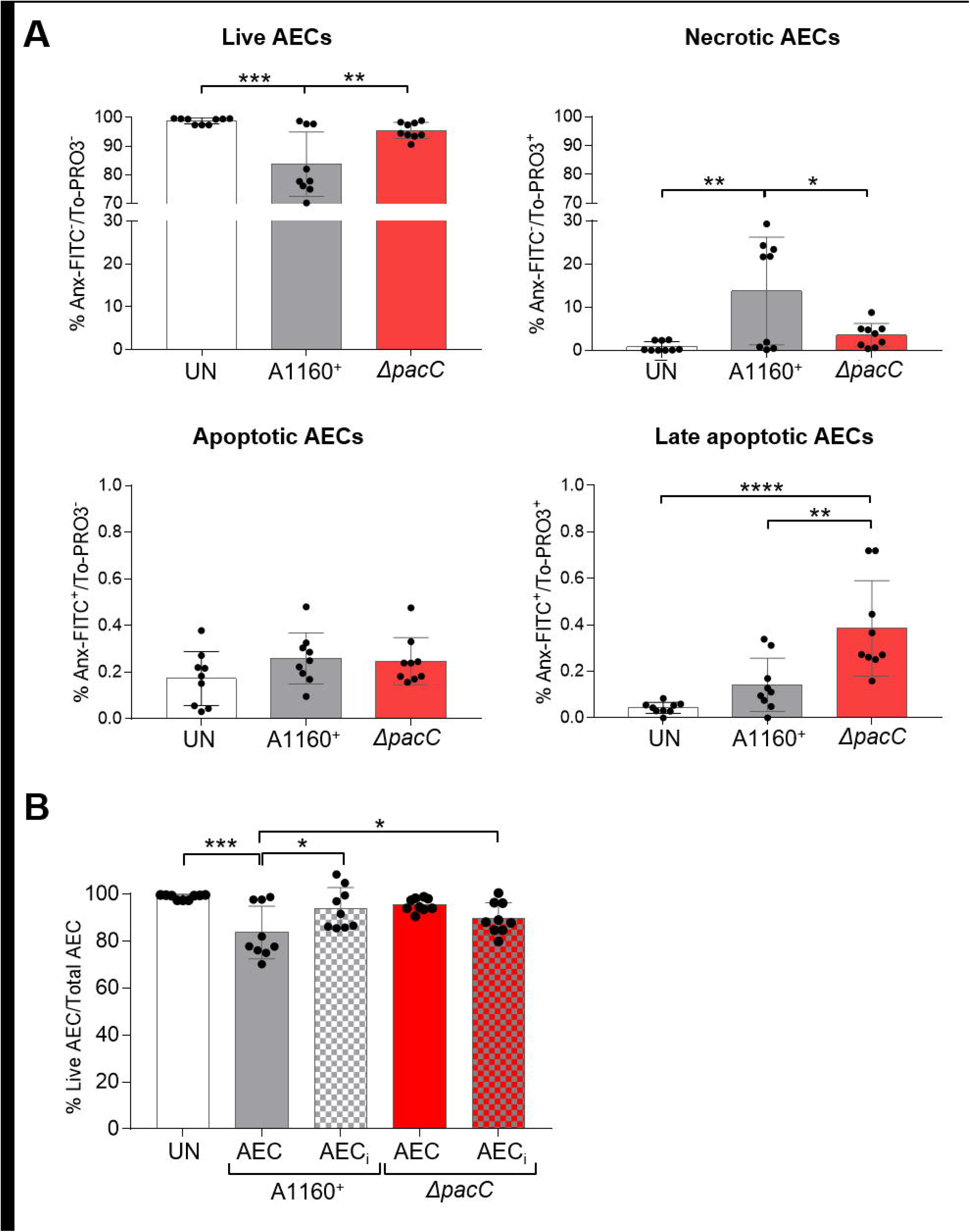
Single-cell analysis of host outcomes upon upon *Af*-AEC interaction indicates that *Af* spore uptake is not the principal cause of AEC death. **(A)** Percentage of live (Anx-FITC^−^/TO-PRO3^−^), necrotic (Anx-FITC^−^/TO-PRO3^+^), apoptotic (Anx-FITC^+^/TO-PRO3^−^) and late apoptotic (Anx-FITC^+^/TO-PRO3^+^) calculated on the total of AECs analysed. **(B)** Percentage of live (Anx-FITC^−^/TO-PRO3^−^) AECs and AEC_i_ calculated on the total of AECs analysed and compared with the percentage of live AECs in uninfected samples. For A-B, infections of A549 monolayers with A1160^+/tdT^ and Δ*pacC*^A1160+/tdT^ were carried out for 16 hours (in biological and technical triplicates) and statistical analysis was carried out using an ordinary 1-way ANOVA with Holm-Sidak’s multiple comparisons. ****p ≤ 0.0001, ***p ≤ 0.001, **p ≤ 0.01, and *p ≤ 0.05

### AECs efficiently kill *Af* spores upon uptake

While viability of internalizing AECs was largely unaffected, our findings using fluorescence microscopy and IFC suggested that *Af* might undergo killing upon uptake. In order to measure *Af* viability upon uptake by AECs, fluorescence-activated cell sorting (FACS) was used to sort pools of 100 AEC_i_, after infection with A1160^+/tdT^ and Δ*pacC*^A1160+/tdT^ for 8 and 16 hours. AEC_i_ pools were subsequently lysed to release the intracellular fungal elements and to perform fungal viability counts **(Fig. 5)**. As we previously demonstrated that at 6 hours post-infection, a higher fungal burden was present in AEC_i_ containing A1160^+/tdT^ compared to the Δ*pacC*^A1160+/tdT^ counterparts **(Fig. 3A)**, the stoichiometry of uptake of the two isolates was determined at 8 and 16 hours post-infection for normalization purposes **(Fig. 5A)**. Similarly, to what observed previously, the number of A1160^+/tdT^ spores contained by 100 AEC_i_ (148.4 ± 18.7 and 128 ± 4.8) was higher than the Δ*pacC*^A1160+/tdT^ counterparts (124.8 ± 6.4 and 115.8 ± 5.7) at both time-points, but the difference between the intracellular fungal burden for A1160^+/tdT^ and Δ*pacC*^A1160+/tdT^ infections was only statistically significant at 8 hours **(Fig. 5A)**. Viability counts of internalised *Af* spores revealed a rapid killing of the intracellular fungal population, whereby by 8 hours post-infection, ~76.2% of the internalised A1160^+/tdT^ fungus was killed by AECs **(Fig. 5B)**. Importantly, fungal killing upon uptake by AECs was almost complete by 16 hour of infection, whereby 90.2% of the internalised A1160^+/tdT^ fungal population was not retrieved in the viable counts **(Fig. 5B)**. Furthermore, despite not reaching statistical significance, AECs were able to curtail even more drastically the growth of the internalised Δ*pacC*^A1160+/tdT^ mutant compared to the respective parental isolate and by 16 hours, 95.0% of the internalised Δ*pacC*^A1160+/tdT^ was killed **(Fig. 5B)**. Taken together, our findings demonstrates a a striking ability of AECs to quell fungal spores upon uptake, strongly suggesting that AECs make a critical contribution to healthy clearance of inhaled *Af* spores by means of their phagocytic activities.

**Fig. 5:**
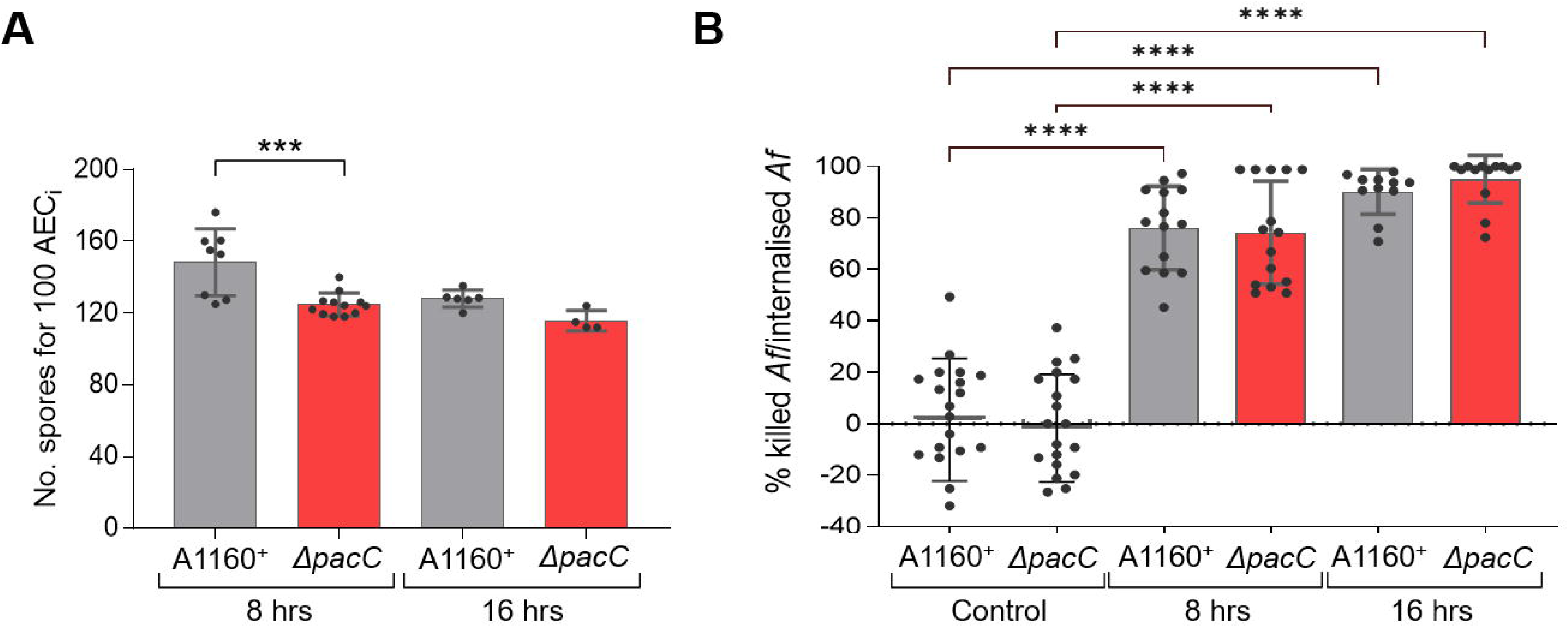
Sorting and lysing of AEC_i_ to perform fungal viable colony count shows rapid and almost complete killing of intracellular *Af*. **(A)** Stoichiometry of A1160^+/tdT^ and Δ*pacC*^A1160+/tdT^ uptake at 8 and 16 hours post-infection of A549 monolayers, expressed as number of *Af* spores internalised by 100 AEC_i_ (n. = 1-5 pools of 100 AEC_i_ for each sample in biological triplicates, one-way ordinary ANOVA and Holm-Sidak’s multiple comparisons test). **(B)** Percentage of dead intracellular *Af* in AEC_i_ relative to internalised *Af* at at 8 and 16 hours post-infection of A549 monolayers with A1160^+/tdT^ and Δ*pacC*^A1160+/tdT^. Significance for each time- and strain-specific sample is shown relative to the respective 100 *Af*^tdT^ spore sorting control (n. = 3-5 pools of 100 AEC_i_ for each sample in biological triplicates, non-parametric 1 non-parametric Kruskal-Wallis test with Dunn’s multiple comparisons test relative to uninfected control). ****p ≤ 0.0001, ***p ≤ 0.001, **p ≤ 0.01, and *p ≤ 0.05

### *Af* is internalized by AECs during infection in mice

While multiple studies have reported spore uptake by immortalised and/or primary AECs in *in vitro* infection systems and *ex vivo* organ culture models [26–32, 35, 36, 45, 46], no compelling evidence of *in vivo* spore uptake by AECs has been published so far. In order to test if *Af* spores are internalised by AECs during mammalian infection, we optimized the isolation and typing of AEC_i_ from mice **(Fig. 6A)**. To maximize the resolution of *Af*-AEC complexes and limit the intereference of phagocytic activities by innate immune cells, an established leukopenic mouse model was used for the experiment [47] and mice were infected with the dtTomato-expressing strain ATCC46645^tdT^ [48] for 8 hours. Following enzymatic digestion of the lung tissue, the multiplex IFC platform was modified and optimized to type epithelial cell subpopulations based on antibody-mediated labelling using the following murine markers: i) EpCam (PE-Cy7-conjugated antibody), ii) Podoplanin (PE-conjugated antibody) and iii) CD74 (FITC-conjugated antibody). While EpCam is routinely used as a generic marker for epithelial cells [49–51], Podoplanin and CD74 have been demonstrated to be eccelent markers to differentiate between alveolar type-I and type-II epithelial cells respectively [52, 53]. Using the experimental pipeline described, we were therefore able to extract and identify both type-I (Podoplanin^+^ CD74^−^ EpCam^+^) and type-II (Podoplanin^−^ CD74^+^ EpCam^+^) alveolar epithelial cells interacting with *Af* in IFC **(Figs. 6A-C)** and FACS-mediated fungal viability assays **(Fig. 6D)**. Alveolar type-I cells cover 95% of the alveolar surface, while alveolar type-II only comprise <5% of the alveolar surface; however, type-II are more numerous and represents 60% of alveolar epithelial cells [54]. Reflecting the actual alveolar composition, the number of type-II alveolar epithelial cells obtained was significantly higher than the number of type-I cells (787 ± 181.5 and 570 ± 103.2 respectively on 2000 EpCam^+^ cells surveyed) **(Fig. 6B)**. Once harvested directly from infected leukopenic mice after 8 hours of infection, respectively 1% and 3.5% of the murine type-I and and type-II alveolar epithelial cells (on 800 cells surveyed) were found to internalise *Af* using IFC **(Fig. 6C)**. At this time point, the stoichiometry of *Af* uptake by murine primary type-II AECs was not found to significantly differ from that one of cultured human type-II AECs (not shown). In order to measure *Af* viability upon uptake by murine primary AECs obtained from infected dissociated lung tissue, FACS was used to sort pools of 100 AEC_i_, after infection with ATCC46645^tdT^ for 4, 8 and 16 hours. AEC_i_ pools were subsequently lysed to release the intracellular fungal elements and to perform fungal viability counts **(Fig. 6D)**. Viability counts of internalised *Af* spores revealed a rapid killing of the intracellular fungal population by AECs, whereby by 4 hours post-infection, ~91.8% and ~69.9% of the internalised fungus was killed by type-1 and type-II AECs respectively **(Fig. 6D)**. Importantly, fungal killing upon uptake by both types of AECs was almost complete by 16 hour of infection, when >97% of the internalised fungal population was not retrieved in the viable counts **(Fig. 6D)**. In conclusion, our innovative single-cell approach, applied directly to infected mouse lungs, has therefore permitted to demonstrate, for the first time, that *Af* spores are internalised and killed by AECs during whole animal infection.

**Fig. 6:**
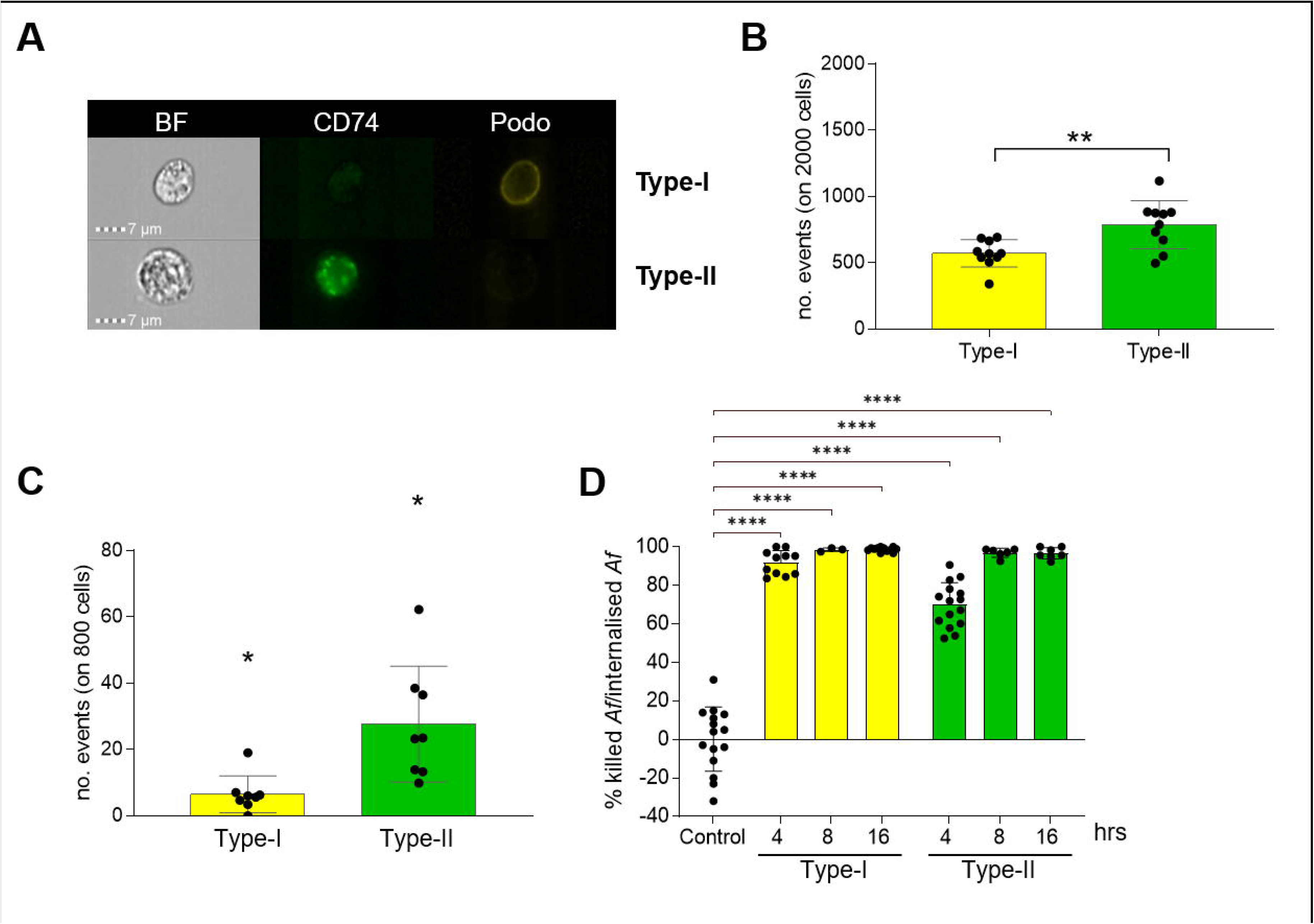
Murine type-II AECs harvested directly from infected leukopenic mice ingest and kill *Af* spores during infection. **(A)** IFC exemplary panels showing type-I (Podoplanin^+^ CD74^-^ EpCam^+^) and type-II (Podoplanin^−^ CD74^+^ EpCam^+^) AECs. **(B)** Quantification of type-I (Podoplanin^+^ CD74^−^ EpCam^+^, yellow) and type-II (Podoplanin^−^ CD74^+^ EpCam^+^, green) alveolar epithelial cells harvested from all murine lungs harvested when surveying 2000 cells. **(C)** Quantification of type-I (Podoplanin^+^ CD74^−^ EpCam^+^, yellow) and type-II (Podoplanin^−^ CD74^+^ EpCam^+^, green) AECs internalising *Af* when surveying 800 cells. Leukopenic mice were infected with 10^8^ spores of ATCC46645^tdT^ for 8 hrs. Following enzymatic dissociation of the lung tissue, antibody-mediated labelling of epithelial (EpCam), alveolar type-I (Podoplanin) and alveolar type-II (CD74) markers was used to identify and quantify epithelial cell subpopulations interacting with *Af*. For B and C, IFC-acquisition was carried out once for the 2 uninfected lungs, while 4 technical replicates were acquired from the 2 pools of 3 lungs obtained from 8-hrs infected mice (unpaired t-test with Welch correction). For C, significance is shown relative to the respective uninfected controls. **(D)** Percentage of dead intracellular *Af* in murine AEC_i_ relative to internalised *Af* at 4, 8 and 16 hours post-infection with ATCC46645^+/tdT^. Significance for each time- and cell-specific sample is shown relative to the respective 100 *Af*^tdT^ spore sorting control (n. = 3-5 pools of 100 AEC_i_ for 3 pools of 5 lungs obtained from 4, 8 and 16 hrs infected mice, non-parametric 1 non-parametric Kruskal-Wallis test with Dunn’s multiple comparisons test relative to uninfected control). ****p ≤ 0.0001, ***p ≤ 0.001, **p ≤ 0.01, and *p ≤ 0.05

### Lung disease significantly impacts uptake of fungal spores

On the premise that epithelial activities provide a potent means of antifungal defence *in vivo* **(Fig. 6)**, we hyphotesised that uptake by AECs would also happen in human primary AECs and that if so, there would likely be some correlations between susceptibility to fungal disease in at-risk patients and aberrancy of this process and its outcomes. We thus tested if commercially available primary human AECs from donors without and with pre-excisting lung disease (asthma and COPD) internalise *Af* spores. IFC enumeration of AECs internalizing the A1160^+/tdT^ isolate demostrated that primary human AECs do indeed internalize *Af* spores after 6 hours of *in vitro* infection **(Fig. 7A)**. Furthermore, COPD-derived AECs were found to internalise significantly more *Af* than their healthy counterpart, whereby the number of COPD-derived AEC_i_ at 6 hours of infection (n = 711.3 ± 249.4 cells on 8000 cells surveyed) was over 4 times higher that the number of healthy-derived AEC_i_ (n = 158.5 ± 119.6 cells on 8000 cells surveyed) **(Fig. 7A)**. As visual examination of the IFC-derived fluorescence microscopy panels confirmed that single AECs are able to internalize multiple *Af* spores, we measured stoichiometry of uptake; however, no significant difference was observed in the number of AEC_i_ containing one or more than one spore in primary human AECs from donors without and with pre-excisting lung disease (not shown). Nevertherless, the fungal burden within all AEC_i_ surveyed (n=8000) was significantly higher for infections of COPD-derived AECs, with 1522 (± 290.9) A1160^+/tdT^ spores internalized by COPD-derived AECs but only 279.3 (± 161.9) spores internalized by healthy AECs **(Fig. 7B)**. Taken together, these results indicated that COPD significantly impacts uptake of fungal spores, as also corroborated when using primary human AECs, extracted, purified and maintained in culture from lung resections sourced locally from the respiratory clinics of the Manchester University NHS Foundation Trust (MFT) via collaboration with the ManARTS Biobank. AECs were derived from two donors with and without COPD and IFC quantification of spore uptake **(Fig. 7C)** substantiated previous observations with commercially available AECs **(Figs. 7A-B)**. Accordingly, COPD-derived AECs internalised significantly more A1160^+/tdT^ spore (n = 1214 ± 271.2 cells for D3 and 958.3 ± 139.7 cells for D4 on 8000 cells surveyed) than healthy AECs (n = 622.6 ± 178.4 cells for D1 and 596.0 ± 189.7 cells for D2 on 8000 cells surveyed) **(Fig. 7C)**.

**Fig. 7:**
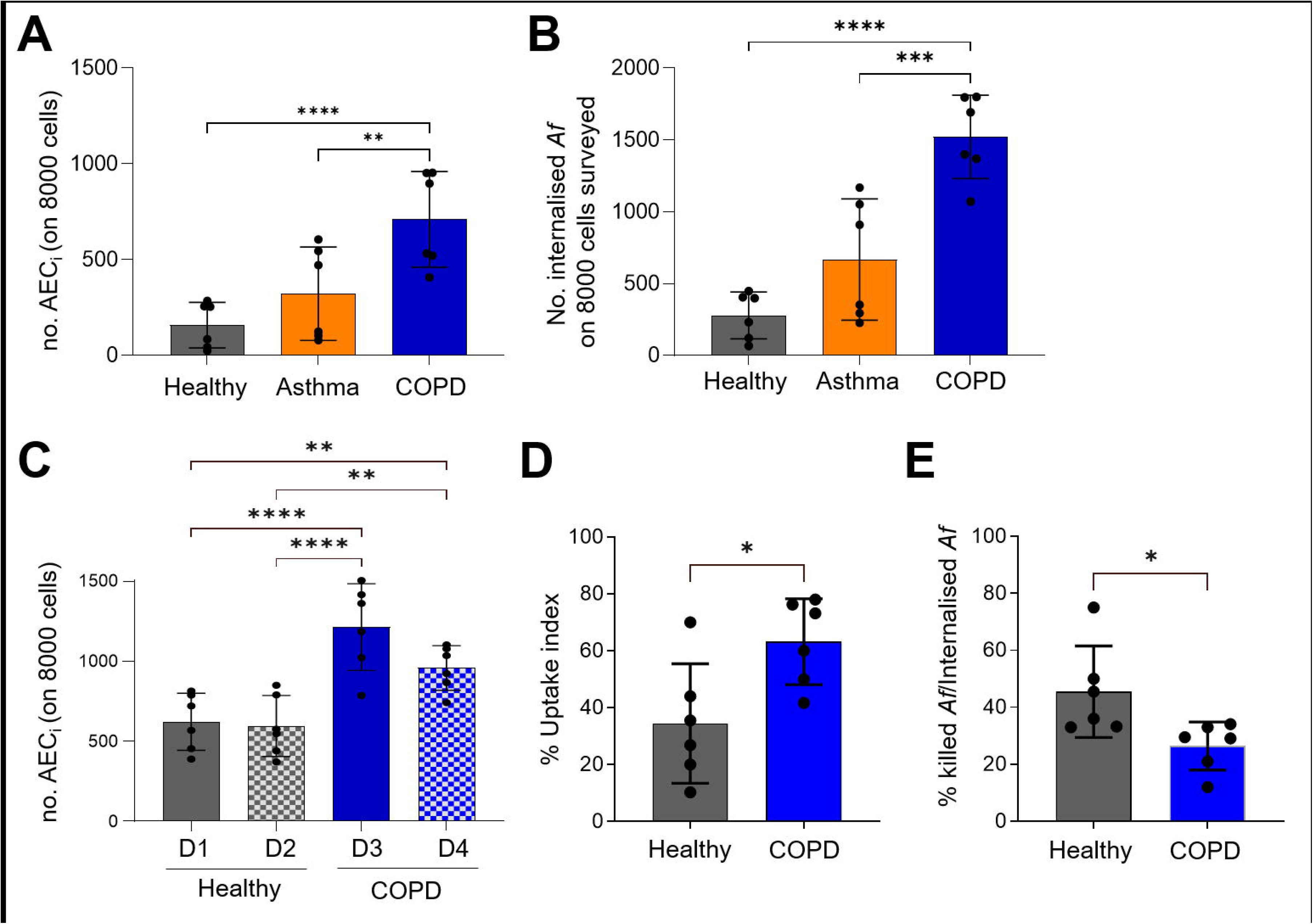
Single-cell comparison of uptake and intracellular killing by primary human AECs indicate that COPD-derived AECs are aberrant in their interaction with fungal spores. **(A)** Quantification of AEC_i_ on 8000 commercially derived cells surveyed by IFC. **(B)** Number of *Af* spores internalised on 8000 AEC surveyed in A by IFC. **(C)** Quantification of AEC_i_ on 8000 locally sourced cells surveyed by IFC. **(D)** Uptake index by and **(E)** percentage of intracellular killing within commercially derived AEC_i_ as quantified by live-cell microfluidic imaging. With the exception of panel C, AEC tested were purchased from Lonza (Healthy = 2547, TAN 29807, Asthma = 2932, TAN 23711 and COPD = 2934, TAN 23484). For C, cells were obtained from locally sourced lung resections obtained via collaboration with the MFT ManARTS Biobank. For A-D, infections of epithelial monolayers with *Af* were carried out for 6 hours, while in E infections were carried out for 20 hrs (in biological duplicates and technical triplicates, ordinary 1-way ANOVA with Holm-Sidak’s multiple comparisons for A-C and unpaired t-test for D-E). ****p ≤ 0.0001, ***p ≤ 0.001, **p ≤ 0.01, and *p ≤ 0.05

Via microfluidic live-cell semi-automated imaging of primary epithelial monolayers exposed to fungal challenge we then determined if intracellular fate of internalized *Af* was also impacted by COPD **(Figs. 7D-E)**. In support of our previous observations **(Figs 7A-C)**, uptake index of COPD-derived AECs after 6 hours of infections was twice as much (63.2 ± 15.1) as that of healthy AECs (34.5 ± 21.0) **(Fig. 7D)**. Most importantly, in agreement with the central hypothesis that the respiratory epithelium plays a critical role in pathogen protection, we demonstrated that in COPD-derived AECs increased spore uptake **(Figs. 7A-D)** correlates with the inability of gorging COPD-derived AECs to quell intracellular *Af* as efficiently as healthy primary AECs **(Fig. 7E)**. After quantification of uptake at 6 hours post-infection, the intracellular fate of internalized A1160^+/mSca^ was followed in real time for further 14 hrs. As mScarlet fluorescence is rapidly quenched upon intracellular killing [55], lack of germination and quenching of mScarlet fluorescence in internalized A1160^+/mSca^ was quantified as a proxy of intracellular killing. Intracellular killing of A1160^+/mSca^ was significantly lower (about half) within COPD-derived AECs (26.5 ± 8.4) than healthy AECs (45.5 ± 16.0) **(Fig. 7E)**. Taken together, this findings suggest that altered AEC responses to fungal challenge may represents a risk factor for aspergillus-related diseases in COPD.

## Discussion

Following inhalation, the respiratory epithelium is the initial point of contact for a multitude of respiratory pathogens of bacterial, viral and fungal origin, which subsequently elicit highly varied and multifaceted host responses. Entry into host mucosae has been conventionally recognised as a pathogenic strategy exploited by microbes to thrive within the host and further perpetuating infection; however, growing evidence demonstrate that the airway epithelium also plays a crucial, and so far underestimated, role in host defence by opsonising, ingesting and quelling inhaled microorganisms [37]. Several published evidence corroborate that uptake by AECs offers protection against infection by different respiratory pathogens of bacterial origin, such as *Pseudomonas aeruginosa*, *Klebsiella pneumoniae* and species of the *Burkholderia cepacia* complex (Bcc) [37]. In these cases, AEC-driven mechanisms of microbial clearance are largely species-specific. While sometimes AECs are directly microbicidal following bacterial uptake (as in the case of Bcc species [56–60] and uncapsulated *K. pneumoniae* [61–63]), in other instances uptake-induced apoptosis and shedding of infected AECs serves as a way of clearing these cells from the airway epithelium (as in the case of *P. aeruginosa* [64–68]). Notably, these microbes have in common the tropism for patients with an impaired immunity or pre-existing lung diseases. In these patients, the lack of innate defences is likely exacerbated by deficient AEC-mediated clearance, thefore highlighting the critical importance of a healthy respiratory niche in delivering efficient defence against inhaled pathogens. *Af* is also an opportunistic pathogen of the human lung; as such, the broad spectrum of diseases caused by this filamentous fungus are largely dependent on impaired (as in patients with acute leukaemia or recipients of allogenic hematopoietic stem cells and solid organ [3–5]) or overzealous (as for asthmatic and cystic fibrosis sufferers [9, 10]) host immune responses or the presence of underlying lung diseases (such as COPD [6, 7]). As most *Af*-related disease manifestations initiate with the inhalation of fungal conidia and eventually lead to the destruction of the pulmonary parenchyma, understanding the AEC-driven mechanisms of microbial clearance is of pivotal importance. We hyphotesised that *Af* uptake and killing by AECs is important for driving efficient fungal clearance *in vivo* and that defective spore uptake and killing would represent major risk factors for aspergillus-related diseases. To this end, we employed state-of-the-art single-cell technologies on *in vitro* and *in vivo* infections to interrogate the complexity of *Af*-AECs interactions and demonstrated that *Af* spore are internalised and killed by immortalised and primary human AECs, and by primary murine AECs during mammalian infection, thus indicating epithelial activities significantly support efficient fungal clearance during mammalian infection.

Using both *in vitro* and *ex vivo* infection models, *Af* uptake has been previously demostrated to be one of the earliest events following spore adhesion to AECs [69]. In these models, approximately 30%–50% of adherent *Af* is internalised by bronchial or type-II alveolar epithelial cells in a concentration- and time-dependent manner [26–28, 30, 70]. However, quantification of uptake by AECs has often relied on indirect methods, such as the nystatin protection assay. These methods anly allows quantification of *Af*, not AECs and most importantly, they rely on fungal viability counts and are therefore unable to accurately separate and quantify uptake from intracellular killing within AECs, as only viable *Af* which has been internalised and survived intracellularly for a set amount of time is eventually enumerated. On the contrary, our direct approaches combine differential fluorescence staining coupled with fluorescence microscopy, IFC [38] or live-cell microfluidic imaging, allowing the direct quantification of both internalised spores **(Fig. 1B)** and internalising AECs **(Fig. 2B)** at the single-cell level and further permitting to establish stoichiometry **(Fig. 1C, Fig. 3 and 5A)** and outcomes of the complex *Af*-AECs interactions at different time-points **(Fig. 4)**. Comparison of uptake indexes *in vitro* showed a rapid time-dependent ~4 fold increase in both the number of internalized spores (relative to total pathogen inoculum) **(Fig. 1B)** and AEC_i_ **(Fig. 2B)** from 2 hours to 6 hours post-infection. Single-cell IFC-based analyses showed that, after infection for 8 hours, while 68.4% (± 15.59%) of the AEC_a_ were attached to germinated *Af,* only 14.5% (± 4.2%) of the AEC_i_ were containing germinated spores **(Fig. 2C-D)**, indicating *Af* might undergo killing upon uptake by AECs.

In *in vitro* infections, epithelial uptake can lead to *Af* killing or, more rarely, intraphagosomal occupancy [33–35, 71–73]. Internalised conidia are quickly trafficked through the endosomal system to the phagolysosomes, as demostrated by the rapid acquisition of late endosomal/lysosomal markers and cathepsin D [29, 72]. Population-scale analyses further indicate that phagosomal acidification results in killing of 97-98% of the internalised conidia within 12-24 hrs, while ∼2-3% of the intracellular *Af* remain viable and in a third of the cases can eventually germinate by 36 h, without lysis of the host cells [29]. Indeed, in our assay, FACS-mediated sorting of AEC_i_ coupled with fungal viability counts revealed a rapid killing of the intracellular *Af* population, whereby ~76.2% and ~90.6% of the internalised fungus was killed within immortalized AECs by 8 and 16 hours post-infection respectively **(Fig. 5B)**. No direct single-cell study on uptake-induced apoptosis or cell death has been so far published in order to understand the relevance of *Af* uptake with regards to AEC survival, but *in vitro* infection assays indicate that *Af* is able to inhibit A549 apoptosis induced by TNF-α, staurosporine and cycloheximide (CHX) [42, 74]. This suggest that pathogen-mediated inhibition of host apoptosis or cell death might facilitate longevity within the intracellular niche. In agreement with our previous results using *in vitro* detachment assays [26], IFC analyses showed that *Af* causes epithelial decay since early-stages of the *Af*-AEC interaction, whereby a significant decrease (15.15%) of live AECs and congruently, a significant increase (12.87%) of necrotic AECs, was measured upon AECs infection with *Af* for 16 hrs relative to uninfected monolayers **(Fig. 4A)**. Importantly however, uptake of fungal spores was ruled out as the major cause of contact-dependent perturbations of epithelial integrity upon *Af* infection *in vitro*, as AEC_i_ did not undergo more cell death than uninfected cells, with 93.98% AEC_i_ still alive at 16 hours post-infection **(Fig. 4B)**. Our findings therefore demonstrated that, while the *Af*-AEC interaction *in vitro* results in disaggregation and damage of the infected monolayers, *Af* spore uptake is not the main cause of the epithelial cell death detected. On the contrary, AECs possess a striking ability to quell fungal spores upon uptake, strongly suggesting that AECs make a critical contribution to healthy clearance of inhaled *Af* spores by uptake.

Suboptimal killing of internalised *Af* spores might result in AECs serving as a fungal reservoir for latent occupation and immune evasion [33–35]. Accordingly, Liu *et al.*, 2016 has recently demonstrated a direct connection between mammalian α5β1 integrin-mediated spore uptake by AECs and polmunary infection [75]. In *in vitro* infection assays, the inhibition of integrin functionality using anti-β1 or −α5 antibodies or siRNA reduces internalisation of *Af* by ~50%, as does the administration of an antibody against the α5β1 integrin fungal cognate, namely the thaumatin-like protein CalA [75]. Importantly, intraperitoneal injection of the anti-CalA antibody prior to *Af* infection in cortisoneacetate-treated mice also increased survival of the infected mice by 20% [75]. However, our study and those of others indicate that AECs potently neutralise the majority of the internalised spores, strongly supporting a curative role for epithelial activities [29, 36, 76]. Spore uptake is dependent on the crucial regulator of host actin dynamics, phospholipase D (PLD), which is activated by exposure to 1,3-glucan on the surface of germinating conidia in a Dectin-1 dependent manner [28]. Accordingly, chemical or siRNA-mediated downregulation of host PLD causes a decrease in *Af* uptake, as does the antibody-mediated blocking of Dectin-1 [26, 28]. Importantly, hematopoietic stem cell transplantees carrying a non-functional truncated version of Dectin-1 have been reported to be at high risk of invasive pulmonary aspergillosis, indicating that Dectin-1 dependent activities of non-hematopoietic cells, such as AECs, participate in curative spore clearance *in vivo*[76]. Furthermore, Chaudhary *et al.*, 2012 reported that bronchial epithelial cells carrying a CFTR mutation (DF508) are impaired in the uptake and intracellular killing of ingested *Af* compared to their wild-type counterparts and also undergo more *Af*-induced apoptosis in response to fungal uptake. Taken together, these evidence suggests that AECs, in collaboration with professional phagoctyes, provide an important, directly microbicidal, defence against *Af* which can become dysfunctional in settings of pre-excisiting respiratory disease or immunosuppression [35, 77].

Pre-existing respiratory disease has been mechanistically correlated to aberrant epithelial uptake of several respiratory pathogens of bacterial origin, leading to the hyphothesis that dysfunctional uptake might contribute to bacterial persistence in certain disease settings [59, 65]. For example, Bcc species are contained in endocytic vesicles and killed within healthy AECs, but in cystic fibrosis (CF) cell lines Bcc-containing endocytic vesicles do not undergo maturation and Bcc species are able to replicate intracellularly causing extensive host damage [56, 57, 59, 60, 78]. Indeed, lung sections of infected CF patients show an abundance of intra-AEC bacteria [79] and Bcc species can be found in CF lungs despite prolonged treatment with antibiotics, indicating a likely intracellular reservoir of Bcc species [80]. Compared to their non-mutated counterparts, cultured human AECs expressing a non-functional CFTR are also defective in the uptake of *P. aeruginosa* and CF patients are hypersusceptibile to lung infections caused by this pathogen [65]. CFTR-mediated uptake of *P. aeruginosa* into healthy AECs results in NF-*κ*B nuclear translocation and eventual apoptosis of the infected host cells [64–68, 81]. Therefore, although not directly microbicidal, *P. aeruginosa* uptake by AECs leads to the desquamation of the infected cells, ultimately driving bacterial clearance and neutralisation of the infection. Accordingly, lung cells from a transgenic CF mice show a decreased ability, relative to wild-type mice, to ingest lipopolysaccharide (LPS)-smooth *P. aeruginosa,* resulting in a significant impairment in bacterial clearance and greater bacterial lung burden[68]. However, AECs expressing a non-functional CFTR undergo apoptosis significantly later than cells expressing wild-type CFTR and in CF murine lungs, no apoptosis is detectable after infection with *P. aeruginosa*, in contrast with their wild-type counterparts [82]. Furthermore, *P. aeruginosa* has an enhanced survival rate in CFTR-deficient cells [83]. As for these bacterial pathogens, spore uptake is likely to range from a useful means of neutralising low-level exposure in health to a driver of invasive growth and pathogenesis [26, 29, 36, 75] where intracellular killing is not possible, as during disease [36].

Our previous *in vitro* findings showed that *Af* non-invasive mutants lacking the transcription factor PacC are unable to elicit early contact-dependent epithelial decay and to secrete soluble effectors of epithelial damage; consequently, they are attenuated for virulence in a leukopenic model of infection [26]. Furthermore, using nystatin protection assays we previously demonstrated that Δ*pacC* mutants are less present inside AECs compared to the respective parental isolates [26], leading us to hypothesise that contact-dependent perturbations of epithelial integrity might in part result from uptake of fungal spores. Using our single-cell approaches, we compared the rate, stoichiometry and outcomes of epithelial uptake of a Δ*pacC* mutant, genetically modified to constitutively express the highly photostable and bright red fluorescent tdTomato [40], to those ones of the respective parental isolate. Comparison of uptake rates by IFC and live-cell microfluidic imaging showed a decrease in Δ*pacC* uptake relative to the parental isolate **(Fig. 3A&D)**; in addition, uptake-stoichiometry analyses for the two strains indicated that the fungal burden within AEC_i_ was significantly higher for the parental isolate than for the Δ*pacC* mutant **(Fig. 3B-D)**. Compared to the parental isolate tested, germination of the Δ*pacC* mutant was significantly impaired within AECs **(Fig. 3E)**. Indipendently of the strain tested, rapid killing of the intracellular fungal populations was observed already at 8 hours post-infection and fungal killing upon uptake by AECs was almost complete by 16 hour of infection, whereby ~90.2% and ~95% of the internalised parental isolate and Δ*pacC* mutant respectively was killed inside AECs **(Fig. 5B)**. Hence, relative to the parental isolate and in agreement with the nystatin protection assay results previously obtained [26], the non-invasive and attenuated Δ*pacC* mutant was less efficiently internalized by AECs and once internalized, more efficiently cleared by these cells. Contrary to what measured for A549 monolayers infected with the parental isolate, infection of A549 cells with the Δ*pacC* mutant did not result in a significant reduction of live AECs in comparison with uninfected monolayers **(Fig. 4A)**, confirming that non-invasive Δ*pacC* mutants lack the ability of causing epithelial decay since early-stages of the *Af*-AEC interaction [26]. Similarly to their counterparts internalizing the parental isolate, AECs internalizing the Δ*pacC* mutant did not suffer more cell death than uninfected monolayers **(Fig. 4B)**, hence excluding the decrease of Δ*pacC* uptake by AECs, as the main reason for the inability of Δ*pacC* mutants to cause epithelial disaggregation and damage *in vitro* and for its attenuation during infection in leukopenic mice [26]. These results point to PacC as crucial in mechanistically mediating *Af* uptake and further intracellular processing by AECs, thereby granting currently ongoing investigations on its role in mediating epithelial activities in response to fungal infection.

A deep understanding of the mode of uptake of *Af* by AECs and the relevance of this process in the context of mammalian disease has been hindered by i) the heterogeneity and complexity of the *Af*-AECs interaction, ii) the limitations of population-scale *in vitro* analysis and iii) the difficulties associated with isolating and culturing primary type-I and −II alveolar cells. While spore uptake has beed reported for immortalized and/or primary AECs in *in vitro* infection systems and *ex vivo* organ culture models [26–29, 31, 32, 36, 45, 46, 75], no compelling published has so far demonstrated *in vivo* spore uptake by AECs. On the contrary, no spore uptake by bronchial epithelial cells of mice has been observed *in vivo* using bioimaging of histological sections combined with transmission electron microscopy (TEM) [84]. However, these findings are derived specifically from the study of bronchial, not alveolar, epithelial cells and, considering the low frequency of spore uptake *in vitro* (and likely *in vivo*), do not exclude that spore internalisation by AECs might occur during mammalian infection [84]. Applying our flow cytometric experimental pipeline for the analysis of *Af*-AECs interaction directly to infected and dissociated murine lungs, we were able, for the first time ever, to demonstrate *Af* spores uptake by AECs and relevance of this process during murine infection. Antibody-mediated typing of AECs subpopulations [52, 53] following enzymatic digestion of infected lung tissue revealed that, after 8 hours of infection, respectively 1% and 3.5% of the murine type-I and and type-II AECs internalise *Af* **(Fig. 6C)**. More importantly, *Af* intracellular viability within murine type-I and and type-II AECs rapidly and drastically declined over time, and only <3% of the internalised fungal population was still viable within both types of AECs by 16 hour of infection **(Fig. 6D)**. To the best of our knowledge, this is the first ever published evidence that, during mammalian infection, *Af* spore uptake occurs, albeith at low rates, and that AECs possess a striking antifungal microbicidal activity, which we hyphotesise can pose a critical contribution to healthy clearance of inhaled *Af* spores.

The discovery that uptake by AECs can provide a potent means of antifungal defence following *Af* exposure during infection in leukopenic mice **(Fig. 6)** opens up many possibilities in terms of understanding human susceptibility to fungal disease and addressing it. Using both commercially available and locally-sourced AECs, we were able to demonstrate not only that primary human AECs internalise and kill *Af* spores, but also that COPD significantly impacts uptake and intracellular killing of *Af* spores by AECs **(Fig. 7)**. COPD-derived AECs were found to internalise significantly more *Af* than their healthy counterpart **(Fig. 7A-D)** more importantly, gorging COPD-derived AECs were unable to quell intracellular *Af* as efficiently as healthy primary AECs **(Fig. 7E)**. *Af* is the most commonly isolated fungus in the sputum of COPD patients, both at steady state and during exacerbations[85] and IA constitutes a significant complication of COPD [7, 8]. Recent estimates indicate that nearly 3.9% of COPD patients admitted to hospital annually develop IA, resulting in 540,451 - 977,082 predicted deaths annually [8]. While bacteria and virus represent the largest causes of exacerbations in COPD patients, accounting for approximately 70% of cases[86], fungal pathogens, such as *Af*, are becoming increasingly recognised as an important etiological cause of exacerbations. Notably, exacerbations due to respiratory infections are the cause of a significant proportion of the morbidity and mortality associated with COPD, which is listed by the World Health Organisation (WHO) as the third leading cause of death in the world, causing over 3 million deaths globally a year. Prevention of exacerbations would thus represent a key strategy for the clinical management of COPD [87–89]; however, our current understanding of the mechanisms underlying the increased susceptibility of these patients to respiratory infections are limited due to the inherent heterogeneity of COPD and the lack of standardised and high-throughput *in vitro* and *in vivo* models of disease [90–93]. The development of a highly controlled *in vitro Af* infection model exploiting commercially available and locally-sourced primary human AECs from healthy donors and COPD patients has allowed us to start deciphering the mechanisms underlying the increased susceptibility of COPD patients to fungal infections. More experiments are currently ongoing to establish donor-dependent variance of spore uptake and intracellular killing by AECs from patients with different COPD severities and if altered protective activities against *Af* in COPD AECs concurs with dysregulation of concomitant immune responses; however, our findings suggest that altered AEC responses to fungal challenge may represents a risk factor for IA in COPD.

Our single-cell approach unmasked the complexity of the *Af*-AECs interactions with respect to the relevance of *Af* uptake by AECs in the context of mammalian disease and susceptibility to fungal infection. Rates, stoichiometry and outcomes of uptake of a non-invasive and attenuated Δ*pacC* mutant and the respective parental isolate were determined, revealing a striking ability of AECs to quell fungal spores upon uptake *in vitro*, during whole animal infection and in healthy human donors. A highly controlled *in vitro Af* infection model exploiting both commercially available and locally-sourced primary human AECs from healthy donors and COPD patients has further allowed us to establish that altered AEC responses to *Af* may represents a risk factor for *Af*-related diseases in COPD. These important findings and powerful technologies are currently driving our further investigations into the molecular, transcriptional and immunological basis of the *Af*-AEC interaction *in vitro*, *in vivo* and in primary AECs to unravel how protective epithelial activities contribute to mucosal tissue homeostasis and pathogen clearance upon *Af* inhalation and can become dysregulated in the settings of pre-excisting respiratory disease, thereby driving *Af*-related diseases. An in-depth understanding of the mechanisms underpinning the antifungal potency of the airway epithelium has major clinical implications as it would aid the identification of immunomodulators to facilitate treatment and limit respiratory damage against respiratory infections caused by *Af*, other fungal and respiratory pathogens.

## Material and methods

### Epithelial cell culture

The epithelial cell line used in this study was the human pulmonary carcinoma epithelial cell line A549 (American type culture collection, CCL-185). A549 cells were maintained at 37°C, 5% CO_2_ in supplemented DMEM (sDMEM, 1% foetal bovine serum, 0.1% penicillin/streptomyn cocktail). Primary human AECs were purchased from Lonza (Healthy = 2547, TAN 29807, Asthma = 2932, TAN 23711 and COPD = 2934, TAN 23484) following isolation from the distal portion of the human respiratory tract in the 1 mm bronchiole area. Alternatively, primary human AECs were isolated and maintained in culture from lung resections obtained via the Manchester Allergy, Respiratory and Thoracic Surgery (ManARTS) Biobank from the Manchester University NHS Foundation Trust (MFT), which is authorised by the National Research Ethics Service (NRES) to release samples to researchers. Both commercially available and locally sourced primary human AECs were maintained at 37°C, 5% CO_2_ in Small Airway Epithelial Cell Growth Medium (Promocell) and used for infections within the ninth passage to avoid senescence. For the microscopy experiments, 10^5^ A549 cells were seeded in 2 ml of sDMEM in 2-well chambered cover glasses (Ibidi) and incubated to ≥ 90 % confluence. For flow cytometric analysis, 10^5^ A549 cells were seeded in 2 ml sDMEM in 6-well plates and incubated to ≥ 90 % confluence. For microfluidic experiments, harvested A549 cells were resuspended in sDMEM at the concentration of 2.5 ×10^6^ cells/ml. CellASIC ONIX M04S-03 Microfluidics plates (Merk) were equilibrated as described by the manufacturers and 10 μl of this suspection was injected in well 6 of the microfluidic plate. The distribution of cells in the wells of the microfluidic plate was checked by microscopy after 5 minutes and if needed, 5 μl of liquid accumulated in well 7 of the plate was removed to favour homogenous distribution of the cells. After 30 minutes from the injection, gravity-driven perfusion was achieved by adding 350 μl and 50 μl respectively to well 1 and 7 of the plate, which was then incubated at 37°C with 5% CO_2_ for 2 days, when cells had reached ≥ 90% confluence.

### Isolation of primary human AECs from lung resections

Lung resections were obtained from the ManARTS Biobank from donors with and without pre-existing lung disease undergoing surgery. Tissues from donors on immune modulatory therapy (for example methotrexate, azathioprine) or with the following diseases were excluded: i) rheumatoid arthritis, ii) other inflammatory diseases (for example Sarcoidosis, Ankylosing spondylitis, Crohns, iii) other autoimmune diseases (for example Inflammatory bowel disease, Multiple sclerosis, Type 1 diabetes), iv) Tuberculosis. Once passed the exclusion criteria, healthy donors were defined as donors if showing i) lung function with Forced Expiratory Volume in first second (FEV1) / Forced Viral Capacity (FVC) >0.7 and FEV1 > 80%, ii) no mention of COPD or emphysema in their hospital records and iii) no diagnosed Type 2 diabetis. COPD donors were classified as definite if i) FEV1/FVC <0.7 and COPD box ticked and ever smoker, ii) FEV1/FVC <0.7 and emphysema on CT scan and ever smoker or iii) FEV1/FVC <0.7, ever smoker and positive symptoms or probable if i) FEV1/FVC <0.7, ever smoker but nil else on proforma. Post-surgery diagnostic analysis of the patient samples and records is used to confirm this. Other investigators may have received tissue from the same subjects.

For the isolation of primary human AECs from lung resections, modifications of the published protocols [94–97] were adopted as follows. Importantly, 3-5 grams of resected lung tissue was finely cut up to be placed into 8 ml of freshly-made digestion buffer, containing 0.4 U/ml Liberase (Sigma) + 160 U/ml DNAse (Sigma) + 0.1 mg/ml Dispase (Sigma). After incubation for 45 minutes at 37°C in a water bath, tissue digestion was stopped by adding 10 ml of cold Stop solution (PBS + 2 mM EDTA + 2% FCS). The digestion lung tissue was filtered serially through a 100 μm, a 70 μm and a 40 μm cell strainer and for each filtration step, the cell strainers were rinsed with 10 ml of Stop solution. The cell suspension obtained was filtered at 400 g, room temperature for 5 minutes. After centrifugation, the supernatant was discarded and the pellet was resuspended in 10 ml of cold 1X Red Blood Cell Lysis buffer (Biolegend). Samples were vortexed and incubated in the dark for 10 minutes at room temperature. 2 ml of cold PBS were added and samples were centrufuges at 700 g, 4°C for 10 minutes. After centrifugation, the supernatant was discarded and the cells were resuspended in 10 ml of adhesion medium for macrophages and fibroblasts (22.5 mL DMEM/F-12, 22.5 mL unsupplemented small airway epithelial cell growth medium, Promocell, 10% mL FBS and 160 U/ml DNase I). Cells were transferred to a T75 tissue culture flask and incubated at 37°C for 60-90 minutes to allow macrophages and fibroblasts to attach to the cell culture flask. This step was repeated twice more and AECs not adhering were gently collected and centrifuged at 300 g for 10 minutes at room temperature. Following centrifugation, the pellets were resuspended in 3 mL DMEM/F-12 and the cell suspension was layered on a 1.040–1.089 g/mL discontinuous Percoll (MP Biomedicals) gradient. Centrifugation at 300 g for 20 minutes at 4°C using a swing out rotor was used to generate an enriched layer of AECs at the interface between the two layers of Percoll gradients. The enriched layer of AECs was transferred to a 15 mL centrifuge tube containing 13 mL HBSS (Sigma) and centrifuged at 300 g for 10 min at room temperature. Following centrifugation, pellets were resuspended in 1 mL of Dynabuffer Isolation Buffer 2 for magnetic bead separation with Dynabeads® CD45 and Dynabeads® CD31 (ThermoFisher) as from manufacturers instructions to remove any possible contamination of macrophages and endothelial cells respectively. Note that, as from manufacturers instruction, for each sample, 50 μl of each Dynabeads® were washed and equilibrated before use using 1 ml of Dynabuffer Isolation buffer 1. After incubation of the cell suspension with the beads for 1 hour at 4°C with gentle tilting and rotation, unbound AECs are transferred for culturing in 5 mls of supplemented Small airway epithelial cell growth medium and maintained in culture for up to 2 months. Cell purity was assessed via flow cytometry, following staining with the follwooing antibodies as from manifacturers specification: EpCAM-PE-Cy7 (Biolegend, 324221) for epithelial cells, Podoplanin-PE (Biolegend, 337003) for alveolar type-I cells and CD74-FITC (Santa Cruz, sc-6262) for alveolar type-II cells[52]. Fluorescence of AECs measured using a BD LSRFortessa system (BD Biosciences) and the BD FACSDIVA 8.0.1 software. The results were analysed using the FlowJo v10 software.

### Fungal strains used in this study

The *Af* strains used in this study were A1160^+^[39], Δ*pacC*^A1160+^[41] and Δ*pksP*^A1160+^ [41]. These strains were genetically modified into A1160^+/dtT^ and Δ*pacC*^A1160+/dtT^ to constitutively express the red fluorescent tdTomato (554/581 nm) [40]. The A1160^+^ was also genetically modified into A1160^+/mSca^ to constitutively express the red fluorescent mScarlet (569/594 nm)[55]. The published dtTomato-expressing strain ATCC46645^tdT^[48], a derivative of the clinical isolate ATCC46645[98], was used for murine infections. As tdTomato has high photostability and low pH sensitivity [40], *Af^tdT^* strains retain fluorescence upon intracellular killing, permitting high-throughput assessment of uptake rates and stoichiometry via IFC [38] and flow-cytometry assisted sorting of *Af^tdT^*-containing AECs for downstream analyses. Fungal killing within AECs is then quantified by sorting and lysing *Af^tdT^*-containing AECs and performing viable counts of the intracellular *Af^tdT^*. In contrast, mScarlet has moderate acid sensitivity and is rapidly quenched upon intracellular killing [55]; therefore, *Af^mScar^* strains represent an ideal tool to trace fungal viability within AECs in real-time.

The strains were cultured at 37°C in *Aspergillus* Complete Media (ACM), when necessary supplemented with pyrithiamine for 3 days [99]. For the preparation of *Af* spore suspensions for infection of A549 monolayers, spores were harvested, filtrated through Miracloth and centrifuged at 3500 rpm for 5 minutes. After counting, a 10^6^ spores/ml suspension was prepared in Dublecco’s Modified Eagle Medium (DMEM, Sigma) supplemented with 10% foetal bovine serum (FBS, Sigma) and 1% penicillin and streptomycin (Sigma). The inoculum concentrations were verified performing viable counts of serial dilutions on ACM plates in duplicate. For both microscopy and flow cytometric analyses, confluent cells were infected with 200 μl of a 10^6^ spore/ml suspension in sDMEM following the replacement of cell media with 2 ml of fresh sDMEM. The infected cells were incubated at 37°C, 5% CO_2_ for the indicated hours.

### Generation and verification of tdTomato- and mScarlet-expressing *A. fumigatus* Strains

The plasmid pSK536 was used for the targeted insertion of the tdTomato fluorophore into the *Af* genome [98]. In pSK536, tdTomato is under the control of the constitutively expressed *A. nidulans* glyceraldehyde-3-phosphate dehydrogenase gene (ANIA_08041) promoter (gpdA) and is targeted at the *Af* region intergenic to the AFUA_3G05360 and AFUA_3G05370, where the terminator region of the *Af* histone 2A locus (his2A^t^) is placed. The ampicillin resistance (bla) and the *A. oryzae* pyrithiamine resistance marker (ptrA) markers are present in the plasmid for selection and maintenance in bacterial and fungal cells respectively. The plasmid pSK536 was also modified into pSK536^mScarlet^ for the targeted insertion of the mScarlet fluorophore into the *Af* genome. mScarlet was amplified from the plasmid pmScarlet_C1 (Addgene) using the oligonucleotides mScarlet1 [CACCGTTTATGGTGAGCAAGGGCGAGGCAGTG] with mScarlet2 [TGGCGTTTCTACTTGTACAGCTCGTCCATGCCGCC]. For replacement of the tdTomato fluorophore with the mScarlet one, the plasmid pSK536 was amplified with the oligonucleotides pSK536_1 [TGTACAAGTAGAAACGCCATGTCTATCTTCGAGTA] and pSK536_2 [CTCACCATAAACGGTGATGTCTGCTCAAGCGG] into the final product pSK536^-tdT^. Assembly of mScarlet and pSK536^-tdT^ into pSK536^mScarlet^ was carried out using the GeneArt technology (ThermoFisher) as from manifacturers’ specifications.

*Af* transformations were performed according to the protocol described in Szewczyk et al. 2006[100]. A1160^+^[39], Δ*pacC*^A1160+^[41] and Δ*pksP*^A1160+^ [41] protoplasts were transformed with circular plasmids and *Af* transformants were selected by supplementing the media with 0.5 ug/ml pyrithiamine [ptrA]. 10^5^ spore/ml of the *Af* transformants were grown in 1 ml of *Aspergillus* Minimal Media (AMM)[99] at 37°C for 6 hours for screening of tdTomato and mScarlet fluorescence via flow cytometry **(Figs. S1A&S2A)**. TdTomato and mScarlet fluorescence of transformants was performed using a BD LSRFortessa system (BD Biosciences) and the BD FACSDIVA 8.0.1 software. The results were analysed using the FlowJo v10 software. The presence and targeted integration to the *his2At* genomic locus of the tdTomato cassette in tdTomato-fluorescent transformants was verified by PRC using the oligonucleotides tdTomato1 [CAGTTCATGTACGGCTCCAA] with tdTomato2 [AGATGGTCTTGAACTCCACCA] and tdTomato3 [GTAACTACGCTCAACGTGTT] with tdTomato4 [CTTCCTGTTGATGGAATGG] respectively (Figs. S1B-C). Copy number integration of the tdTomato cassette was checked by Southern analysis using a prtA-specific hybridisation probe generated using the oligonucleotides PtrA_SB1 [GGATAGGGGCGAACTTGAACT] and PtrA_SB2 [TTTGGCTGGACTCTCACAAT] **(Figs. S1B-C)**. Genomic DNA for Southern blotting anlyses was digested with EcoRI. The presence to the *his2At* genomic locus of the mScarlet cassette in mScarlet-fluorescent transformants was verified by PRC using the oligonucleotides mScarlet1 [TCCCCTCAGTTCATGTACGG] and mScarlet 2 [CTTGTACAGCTCGTCCATGC] **(Figs. S2B-C)**. Targeted integration of a single copy of the mScarlet cassette to the *his2At* genomic locus was checked by Southern analysis using *his2A*- and *prtA*-specific hybridisation probes, generated using the oligonucleotides His2A1 [TTCACCTGATTCAGCTGATTG] with His2A2 [TTCACCTGATTCAGCTGATTG] and PtrA_SB1 [GGATAGGGGCGAACTTGAACT] with PtrA_SB2 [TTTGGCTGGACTCTCACAAT] **(Figs. S2B-C)**. Genomic DNA for Southern blotting anlyses was digested with EcoRI.

### Differential fluorescence assay for quantification of internalisation

Following infection, the supernatant media was removed and monolayers were washed three times with warm PBS (PBS, Life Technologies). Infected cells were stained with 1 ml of sDMEM containing 10 mg/ml Concanavalin A-FITC (Con-A, FITC 495/519 nm, Sigma) for 30 minutes at 37°C, 5% CO_2_. For the final 5 minutes of the staining period, 1 ml of sDMEM containing 0.8 mg/ml calcofluor-white (CW, 365/435 nm, Sigma) was added. The stains were removed after incubation and replaced with 2 ml warm PBS. Live cell imaging was performed using a Nikon Eclipse TE2000-E microscope with DIC optics and a 60X (1.20 NA) Plan Apo objective, images were captured with an ORCA-ER CCD camera (Hamamatsu, Welwyn Garden City, UK) driven by the MetaMorph NX 1.1 software for image acquisition. For calcofluor white, a Nikon UV-2A filter cube (excitation filter 355/15 nm BP, dichoric mirror 400 nm LP, emission filter 420 nm LP) was used. For Concanavalin-A-FITC, a Nikon B-2A filter cube (excitation filter 470/20 nm BP, dichroic mirror 500 nm LP, emission filter 515 nm LP) was used. For tdTomato, a Nikon G-2A filter cube (excitation filter 535/50 nm BP, dichroic mirror 565 nm LP, emission filter 590 LP) was used. Images were processed and analysed using the software ImageJ[101]. Images show maximum intensity projections with inverted look-up tables (LUTs). Multiple fields of view (n = 20-42) were pictured at random using 2 well plates of seeded A549 cells which were infected with tdTomato^ATCC^ for the indicated time and stained with ConA-FITC and CW. Intracellular and extracellular *Af* were enumerated and a correction factor was applied to normalise for the experimental variation in the multiplicity of infection (MOI, the ratio of the number of pathogens to the number of cells in each field of view). The MOI for each field of view in each replicate was quantified and averaged across biological replicates. The normalised MOI was then used to correct the internalisation index as follows.

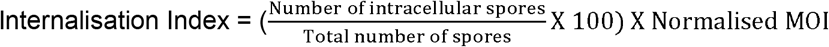

### IFC analysis of *Af*-AEC interaction

The detailed protocol for single-cell analyses of fungal uptake in cultured AECs is decribed in Bertuzzi & Howell, 2021 [38]. Following infection, the *Af* inoculum concentrations were verified performing viable counts of serial dilutions on ACM plates in duplicate. ACM plates were incubated for 24-48 hours at 37°C and colonies were enumerated to calculate the actual inoculum for infection. The deviance of the actual inoculum from the expected inoculum (10^6^ spore/mL) for each replicate determine the coefficient of infection for normalisation purposes. For the preparation of the samples for IFC, the supernatant from infected cells was collected. Infected cells were washed once with pre-warmed PBS and the wash was pooled together with the supernatants from the infected cells. Infected cells were detached from the 6-well plates by incubation for 5 minutes in 1 ml of trypsin-EDTA at 37°C, 5% CO_2_. After 5 minutes, 1 ml of fresh sDMEM was added and detached cells were collected and pooled together with the supernatant and wash. Cells were centrifuged at 1800 rpm for 3 minutes. Pellets were resuspended in 500 μl of 1X Binding Buffer containing 5 μl of Annexin V-FITC (495/519 nm, Biovision), 1 nM TO-PRO™-3 Iodide (642/661 nm, Thermofisher) and 8 ng/ml calcofluor-white (CW, 365/435 nm, Sigma). Samples were incubated in the dark at room temperature for 5 minutes and then centrifuged at 2400 rpm for 3 minutes. As described in Bertuzzi & Howell, 2021 [38], controls to determine the gating strategy included A549 cells i) infected with A1160^+^ with and without staining with CW, ii) pre-treated for 1 hour with the endocytosis inhibitor Cytochalasin D (0.2 μM) with and without infection with A1160^+/dtT^ and iii) cells treated with the apoptosis inducer Staurosporine (1 μM) with and without infection with A1160^+/dtT^. Pellets were resuspended in 1 ml of PBS and then centrifuged at 2400 rpm for 3 minutes. Pellets were resuspended in 50 μl of PBS for IFC processing. Samples were processed using an ImageStream®X Mark II Imaging Flow Cytometer (Merk) with the following settings: Laser 488 at 200V, Laser 561 at 200V, Laser 405 at 75V, Laser 642 at 100V and SSC 2. Total events were filtered in order to exclude clumps of cells or cells out of focus and collected at the number indicated in each experiment. Results were analysed using the IDEAS® software and further gating was performed based on the bright-field size and fluorescent intensity on channel 2 (FITC signal, 480-560), channel 3 (tdTomato signal, 560-595), channel 7 (CW signal, 435-505) and channel 11 (TO-PRO™-3 signal, 642-745). For each sample, as indicated in the result section, 2 or 3 acquisitions of 2000 or 8000 single cells in focus were performed on the ImageStream for technical replicates and each experiment was performed in biological duplicate or triplicates. Data analysis using the IDEAS® software and normalisation of *Af*-AEC interaction and outcomes thereof was performed according to the pipeline described in Bertuzzi & Howell, 2021 in order to normalise across experimental replicates with background subtraction and correction based on the coefficient of infection determined from the viable counts[38]. *Af* germination on, or within, AECs and spore stoichiometry following uptake was enumerated by viable examination of every AEC_a_ and AEC_i_ using the IDEAS® software with biological and technical replicates as indicated in the result section. Timepoint- and strain-specific stoichiometry coefficient was calculated by averaging the number of spores internalised by 100 AEC_i_ in all the replicates (technical and biological) at the time-point of *Af*-AEC interaction indicated.

### Fluorescence-activated cell sorting (FACS) analysis coupled with viability counts to measure *Af* viability within AECs

For the preparation of the samples for FACS, the supernatant from infected cells was collected. Infected cells were washed once with pre-warmed PBS and the wash was pooled together with the supernatants from the infected cells. Infected cells were detached from the 6-well plates by incubation for 5 minutes in 1 ml of trypsin-EDTA at 37°C, 5% CO_2_. After 5 minutes, 1 ml of fresh sDMEM was added and detached cells were collected and pooled together with the supernatant and wash. Cells were centrifuged at 1800 rpm for 3 minutes. Pellets were resuspended in 500 μl of PBS containing 8 ng/ml calcofluor-white. Samples were incubated in the dark at room temperature for 5 minutes and then centrifuged at 2400 rpm for 3 minutes. Pellets were resuspended in 1 ml of PBS and then centrifuged at 2400 rpm for 3 minutes. Pellets were resuspended in 50 μl of PBS for FACS processing using a BD Influx Cell Sorter. Total events were filtered in order to exclude clumps of cells and plotted based on the fluorescent intensity of tdTomato and CW signal. For each strain and time-point indicated, pools (n = 3-5) of 100 AECi (tdTomato^+^/CW^-^) were sorted in sterile H_2_O. AECi were lysed by vigourous votexing and internalised viable *Af* was enumerated following plating of the lysates onto ACM plates and incubation of the plates for 24-48 hours at 37°C. The experiment was performed in biological triplicates. As sorting control, for each biological replicate, 5 pools of 100 *Af*^tdT^ spores were sorted in sterile H_2_O and plated in ACM for culturing and viable counts. In order to calculate the percentage of killing of intracellular *Af* in AEC_i_ relative to internalised *Af*, data normalisation was performed based on the relative time- and strain-specific stoichiometry coefficient calculated previously. Statistical analysis was carried out by comparing each time- and strain-specific time point in the analysis with the respective 100 *Af*^tdT^ spore sorting control.

### Microfluidic live-cell imaging

For microfluidic experiments, *Af* spores were resuspended at a concentration of 10^5^ spore/ml and CW was diluted to 4 μg/ml suspension, both in Fluorobrite sDMEM (ThermoFisher). Plates were prepared as from manifacturer’s instructions and 200 μl of FluoroBrite, 200 μl of *Af* spore suspension, 200 μl of FluoroBrite, and 200 μl of 4 μg/ml CW suspension were added to wells 2, 3, 4 and 5 respectively. *Af* spores were injected onto the A549 monolayers using the Cellasic manifold and the ONIX software (v.5.0) at 37°C, 5% CO_2_. Infection was performed by flowing first fresh FluoroBrite media (from well 2) at 2 psi for 10 minutes and then alternating pulses of *Af* spores (from well 3) at 4 psi (30 seconds), 8 psi (30 seconds) and 0.25 psi (2 minutes) for 3 cycles. After achieving a satisfactory spore distribution, as visualised using a 20x/0.75NA objective and a Nikon TiS Eclipse microscope, the injection of spores was stopped and plates were incubated for 6 hours at 37°C, 5% CO_2_. After 6 hours of incubation, the CW suspension (from well 5) was injected onto the monolayers at 2psi for 10 minutes, followed by a wash with Fluorobrite (from well 2) at 4 psi for 10 minutes. For the following 14 hours of infection, FluoroBrite was set to flow into the chamber at 0.5 psi. These processes were automatically controlled by ONIX software (v.5.0). The 4D multipoint stacked imaging was obtained using a Nikon TiS Eclipse microscope with a 20x/0.75 objective lens, whereby live cell images were taken at two hours intervals over a duration of 14 hours from the injection of CW at 6 hours post-infection. An Hamamtsu Flash 4.0 sCMOS camera with a multiband Semrock filter and the Nikon Elements V.4.2 acquisition software was used to capture and collect emitted fluorescence from CW and TdTomato. Using the Fiji counter[102], the number of internalized spores (tdTomato^+^, CW^−^) and extracellular spores (tdTomato^+^, CW^+^) was enumerated using the 6 hr 3D multipoint image and uptake index was calculated as follow:

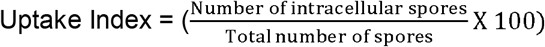

Once identified intracellular *Af* spores at 6 hour post-infection, their growth rate was assessed by measuring the length of the growing hyphae after 14 hours, using Fiji [102].

### Murine infection and lung processing and staining for flow cytometry

Murine infections were performed under UK Home Office Project Licence PDF8402B7 in dedicated facilities at the University of Manchester. *Af* spores were harvested and prepared as previously described[47] and viable counts from administered inocula were determined, following serial dilution, by growth for 24–48 hours at 37°C on ACM. Individually vented cages were used to house the mice, which were anaesthetized by halothane inhalation and infected by intranasal instillation of 40 μl of ATCC46645^tdT^ suspensions at the concentration of 2.5×10^9^ spores/ml (dosage = 10^8^ spores). Leukopenic mice were immunosuppressed by administration of cyclophosphamide (150 mg/kg, intraperitoneal) on days −3, −1, +2 and every subsequent third day, and a single subcutaneous dose of hydrocortisone acetate (112.5 mg/kg) administered on day −1. Mice were culled after 4, 8 and 16 hours of infection and lungs were perfused by injection of 2 ml of dispase solution (0.1 mg/ml Dispase, Sigma) in HANKS buffer (Sigma) directly into trachea. Whole lungs (with trachea and heart) were collected and tissue digestion was carried out at low speed (100 rpm) for 1 hr at 37°C. Hearts and tracheas were removed and lung tissues were minced using sterile scissors prior to addition of 2 mls of digestion buffer containing 0.1 mg/ml Dispase, 0.4 U/ml Liberase TL (Sigma) and 160 U/ml DNase. The samples were incubated for further 10 minutes, 100 rpm, at 37°C. Sample volume was adjusted to 10 ml via the addition of PBS + 2% FBS + 2 mM EDTA. Lungs from 6 infected mice were pooled together in 2 groups of 3, while the lungs from the 2 uninfected animals were processed separately. Lung tissues were filtered through 70 μm and 40 μm strainers and the volume was adjusted to 50 ml via the addition of PBS + 2% FBS + 2 mM EDTA. Samples were centrifuged 1800 rpm for 6 minutes at 4°C and pellets were resuspended n 3 mL RBC lysis for 4 minutes (Sigma). Samples were centrifuged 1800 rpm for 6 minutes at 4°C and the pellets were resuspended in 200 μl of PBS + 2% FBS + 2 mM EDTA. 5 μg/ml of anti-CD16/32 antibody (EBioscience) were added and samples were incubated for 20 minutes on ice and in the dark. Samples were centrifuged 1800 rpm for 6 minutes at 4°C and the pellets were resuspended in 100 μl of staining mix, containing 1 μg/ml of the antibodies anti-mouse EpCam (PE-Cy7, Biolegend), anti-mouse Podoplanin (PE, eBioscience) and anti-mouse CD74 (FITC, Santa Cruz) in 100 μl of PBS + 2% FBS + 2 mM EDTA. Following incubation for 25 minutes in the dark and on ice, 1 nM TO-PRO™-3 Iodide (642/661 nm, Thermofisher) and 8 ng/ml calcofluor-white (CW, 365/435 nm, Sigma) was added and the samples were incubated for further 5 minutes in the dark and on ice. Samples were centrifuged 1800 rpm for 6 minutes at 4°C and the pellets were resuspended in 100 μl PBS for IFC or FACS processing. In parallel, single stain-controls were carried out on a processed lung from an uninfected mouse to determine the gating strategy. For IFC, samples were processed using an ImageStream®X Mark II Imaging Flow Cytometer (Merk) as previously specified for *in vitro* experiments. For each sample (1 uninfected and two 8 hours post-infection), 5 acquisition of 2000 EpCam^+^ single cells in focus were taken as technical replicates. Data analysis was performed using the IDEAS® software and the pipeline previously described. For FACS, samples were processed as previously specified for *in vitro* experiments. Moreover, for each cell type (alveolar type-I Podoplanin^+^ CD74^-^ EpCam^+^ and alveolar type-II Podoplanin^-^ CD74^+^ EpCam^+^)[52] and time-point indicated, pools (n = 1-5) of 100 AEC_i_ (tdTomato^+^ /CW^-^) were sorted in sterile H_2_O and internalised viable *Af* was enumerated and analysed as previously specified for *in vitro* experiments.

### Statistical analysis of data

GraphPad Prism was used to interpret data and p values were calculated through unpaired t tests (with Welch correction as indicated), ordinary 1-way ANOVA with Holm-Sidak’s multiple comparisons test or non-parametric Kruskal-Wallis test with Dunn’s multiple comparisons test, as indicated. Error bars show the Standard Deviation (SD). ****p ≤ 0.0001, ***p ≤ 0.001, **p ≤ 0.01, and *p ≤ 0.05

## ACKNOWLEDGMENTS

A special thanks goes to Caitlin Sergienko, Lea Gregson and Saif Abraham for their technical assistance. This study is independent research supported by the North West Lung Centre Charity at Manchester University NHS Foundation Trust. The views expressed in this publication are those of the authors and not necessarily those of the NHS, the North West Lung Centre Charity or the Department of Health. The authors would like to acknowledge the Manchester Allergy, Respiratory and Thoracic Surgery Biobank and the North West Lung Centre Charity for supporting this project. In addition we would to thank the study participants for their contribution.

## Supplementary Figures

**Fig. S1:**
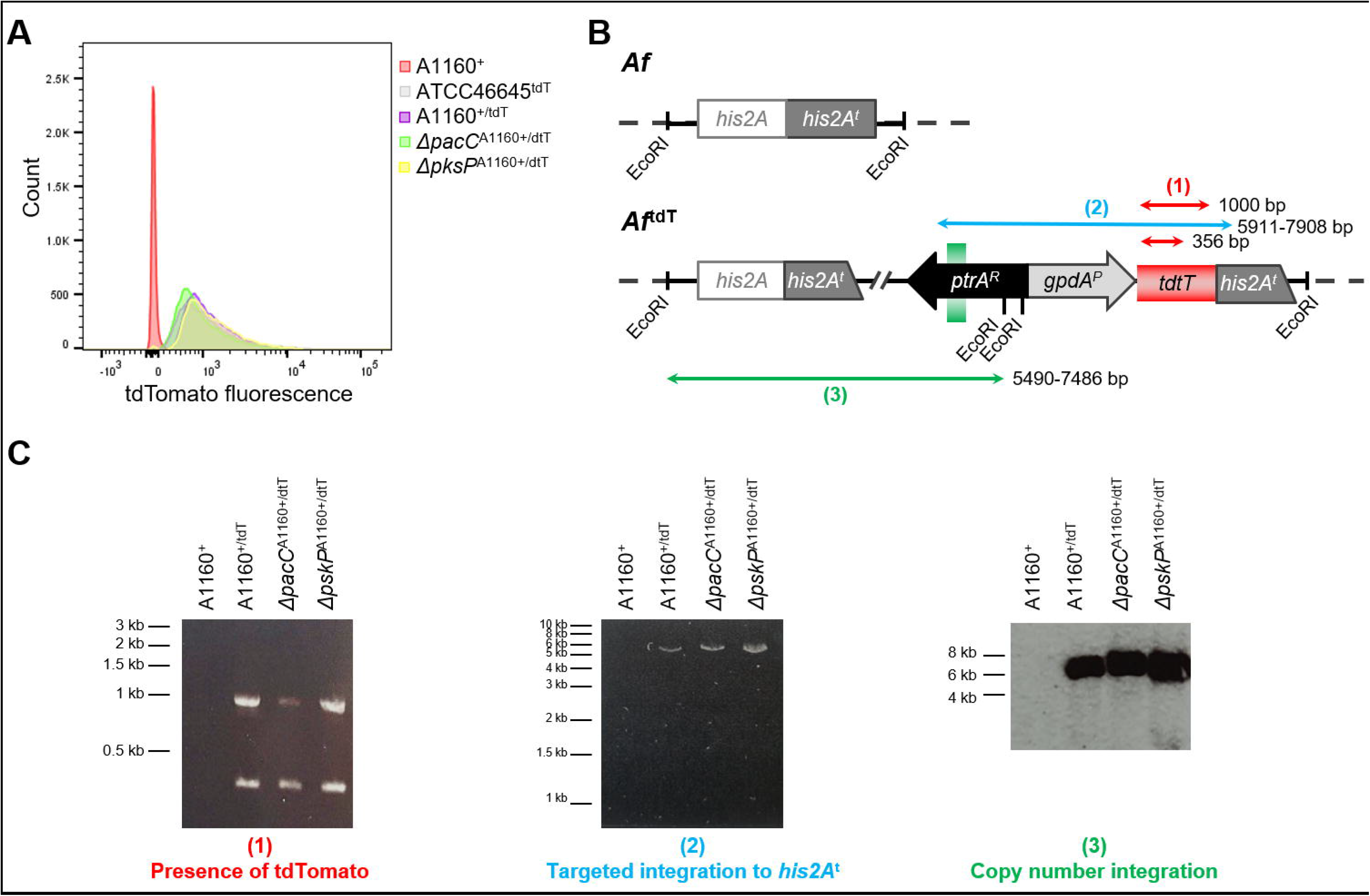
Design and construction of the tdTomato-expressing *Af* strains (*Af*^tdT^). **(A)** Histogram displaying the number of events relative to the tdTomato fluorescence (561 586_15 laser) for the screening of *Af* transformants using flow cytometry. The compararison shows the parental isolate A1160^+^, the published tdtomato-expressing ATCC46645 isolate[48] and a rapresentative A1160^+/dtT^, Δ*pacC*^A1160+/dtT^ and Δ*pskP*^A1160+/dtT^ transformants. **(B)** PCR and Southern blotting strategy for the verification of single, targeted integration of the tdTomato expression construct in A1160^+^, Δ*pacC*^A1160+^ and Δ*pskP*^A1160+/dtT^ transformants. The presence (1) of the tdTomato cassette was verified using the oligonucleotides tdTomato1 and tdTomato2. Two PCR bands are expected from tdTomato-expressing clones (1000 and 356 bp), while no band is expected for the parental isolates A1160^+^. The targeted integration (2) of the tdTomato cassette was verified using the oligonucleotides tdTomato3 and tdTomato4. A single PCR band is expected from tdTomato-expressing clones (5911-7908 bp), while no band is expected for the parental isolates A1160^+^. Copy number insertion (3) of the tdTomato cassette was verified by Southern blotting using a *prtA*-specific hybridisation probe generated with the oligonucleotides PtrA_SB1 and PtrA_SB2. Genomic DNA was digested with EcoRI. No band is expected for the parental isolate, whereas a single band, ranging from 5490 to 7486 bp dependent upon the precise site of integration, is expected for the reporter strain. **(C)** PCR and Southern blotting for the verification of single, targeted integration of the tdTomato expression construct in rapresentative A1160^+/dtT^, Δ*pacC*^A1160+/dtT^ and Δ*pskP*^A1160+/dtT^ transformants.

**Fig. S2:**
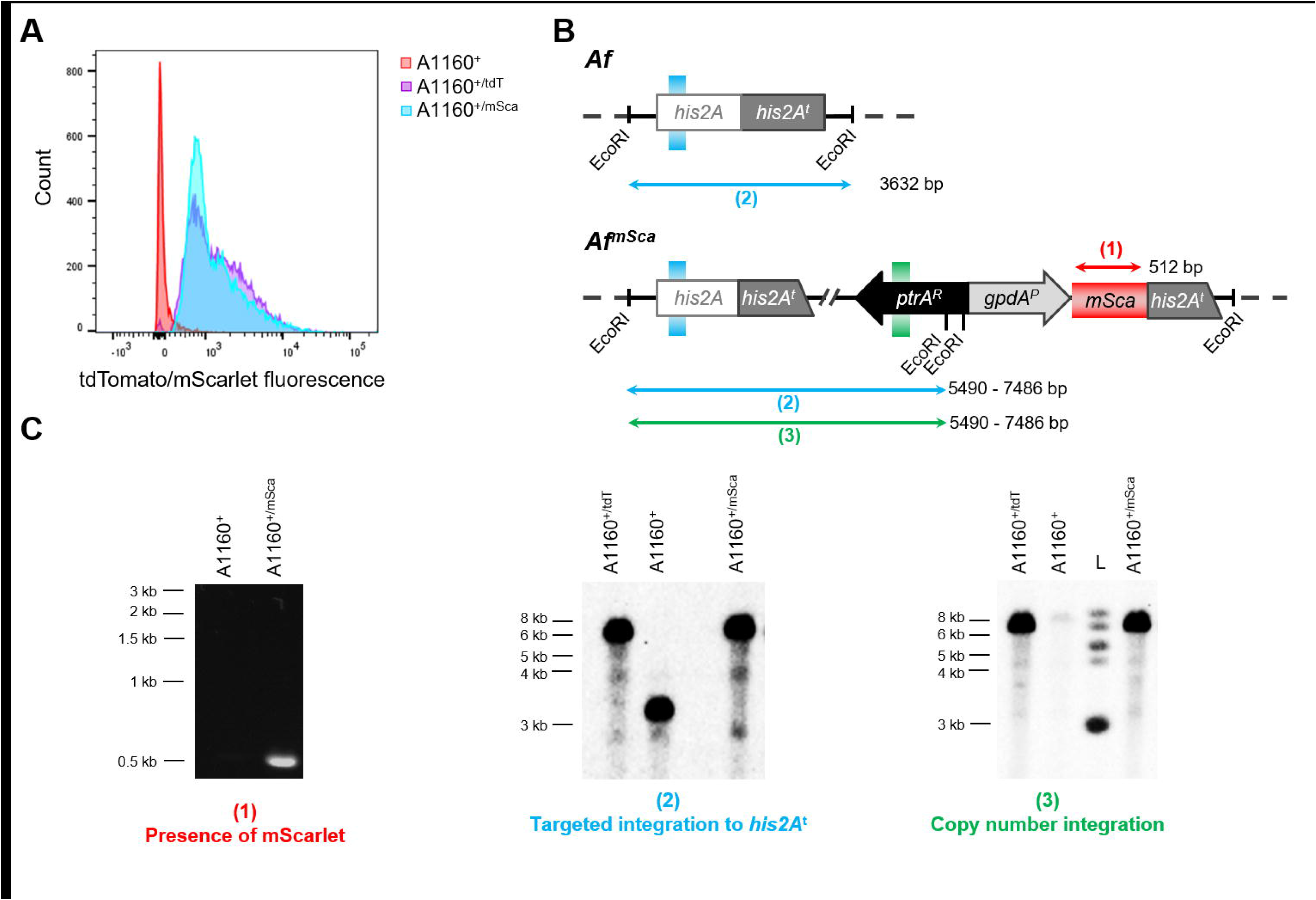
Design and construction of the mScarlet-expressing *Af* strain (*Af*^mSca^). **(A)** Histogram displaying the number of events relative to the mScarlet fluorescence (561 586_15 laser) for the screening of *Af* transformants using flow cytometry. The compararison shows the parental isolate A1160^+^, the rapresentative A1160^+/dtT^ from Fig. S1 and a rapresentative A1160^+/mSca^. **(B)** PCR and Southern blotting strategy for the verification of single, targeted integration of the mScarlet expression construct in A1160^+^. The presence (1) of the mScarlet cassette was verified using the oligonucleotides mScarlet1 and mScarlet2. A PCR band of 512 bp is expected from mScarlet-expressing clones, while no band is expected for the parental isolates A1160^+^. The targeted (2) and single (3) integration of the mScarlet cassette was verified by Southern blotting using *his2A*- and *prtA*-specific hybridisation probes, generated using the oligonucleotides His2A1 with His2A2 and PtrA_SB1 with PtrA_SB2. Genomic DNA was digested with EcoRI. Using the *his2A*-specific hybridisation probe (2), a single band of 3632 bp is expected for parental isolates, while a single band, ranging from 5490 to 7486 bp dependent upon the precise site of integration, is expected for the reporter strain. Using the *prtA*-specific hybridisation probe (3), no band is expected for the parental isolate, whereas a single band, ranging from 5490 to 7486 bp dependent upon the precise site of integration, is expected for the reporter strain. **(C)** PCR and Southern blotting for the verification of single, targeted integration of the mScarlet expression construct in A1160^+/msca^. L = protein ladder

